# TASOR is a pseudo-PARP that directs HUSH complex assembly and epigenetic transposon control

**DOI:** 10.1101/2020.03.09.974832

**Authors:** Christopher H. Douse, Iva A. Tchasovnikarova, Richard T. Timms, Anna V. Protasio, Marta Seczynska, Daniil M. Prigozhin, Anna Albecka, Jane Wagstaff, James C. Williamson, Stefan M.V. Freund, Paul J. Lehner, Yorgo Modis

## Abstract

**Summary:** The Human Silencing Hub (HUSH) complex epigenetically represses retroviruses, transposons and genes in vertebrates. HUSH therefore maintains genome integrity and is central in the interplay between intrinsic immunity, transposable elements and transcriptional regulation. Comprising three subunits – TASOR, MPP8 and Periphilin – HUSH regulates SETDB1-dependent deposition of the transcriptionally repressive epigenetic mark H3K9me3 and recruits MORC2 to modify local chromatin structure. However the mechanistic roles of each HUSH subunit remain undetermined. Here we show that TASOR lies at the heart of HUSH, providing a platform for assembling the other subunits. Targeted epigenomic profiling supports the model that TASOR binds and regulates H3K9me3 specifically over LINE-1 repeats and other repetitive exons in transcribed genes. We find TASOR associates with several components of the nuclear RNA processing machinery and its modular domain architecture bears striking similarities to that of Chp1, the central component of the yeast RNA-induced transcriptional silencing (RITS) complex. Together these observations suggest that an RNA intermediate may be important for HUSH activity. We identify the TASOR domains necessary for HUSH assembly and transgene repression. Structural and genomic analyses reveal that TASOR contains a poly-ADP ribose polymerase (PARP) domain dispensable for assembly and chromatin localization, but critical for epigenetic regulation of target elements. This domain contains a degenerated and obstructed active site and has hence lost catalytic activity. Together our data demonstrate that TASOR is a pseudo-PARP critical for HUSH complex assembly and H3K9me3 deposition over its genomic targets.

## Introduction

Post-translational modification of histones and other chromatin proteins is a central mechanism by which eukaryotic cells regulate chromatin architecture and tune the dynamics of DNA-templated processes. One conserved example is trimethylation of histone H3 lysine 9 (H3K9me3), an epigenetic mark typically associated with low levels of transcription (Kouzarides, 2007). H3K9me3 marks repetitive regions of eukaryotic chromosomes where it presents a binding site for heterochromatin protein 1 (HP1) (Nielsen et al., 2002). HP1 undergoes liquid-liquid phase separation to form a chromatin compartment that somehow excludes RNA polymerase from the DNA (Strom et al., 2017). However, H3K9me3 is also present in transcriptionally-active euchromatin (Becker et al., 2017) – for example over the bodies of certain protein-coding genes (Blahnik et al., 2011). The mark is therefore central in maintaining genome stability and controlling transcriptional programs. In mammals its importance at an organismal level is underlined by recent observations that dynamic regulation of H3K9me3 – catalyzed by multiple lysine N-methyltransferases – is critical for murine development (Nicetto et al., 2019).

Genetic experiments studying position-effect variegation (PEV) in model organisms have identified much of the machinery involved in the formation of H3K9me3 domains (Elgin and Reuter, 2013). Such position effects refer to the influence of the chromatin environment on gene expression. Forward genetic screens for mutations disrupting PEV in *Drosophila* revealed conserved factors required for heterochromatin formation including HP1 itself (Schotta et al., 2002). Analogously, a mutagenic screen in a mouse line with a transgene reporter displaying variegated expression identified numerous epigenetic regulators, the ‘Modifiers of murine metastable epialleles’ (*Mommes*), several of which are specific to mammals (Blewitt et al., 2005; Daxinger et al., 2013). A forward genetic screen that we conducted previously with an integrating lentiviral reporter identified the Human Silencing Hub (HUSH) as a novel regulator of PEV in human cells (Tchasovnikarova et al., 2015). HUSH is a complex of three proteins: Transgene Activation Suppressor (TASOR), M-phase phosphoprotein 8 (MPP8) and Periphilin (PPHLN1). The activity of TASOR is critical in early development: the homozygous mutation L130P, at a conserved leucine in mouse TASOR (identified as MommeD6), is lethal in embryos before gastrulation (Harten et al., 2014). The HUSH complex recruits the H3K9 methyltransferase SET domain bifurcated 1 (SETDB1) to deposit H3K9me3 (Tchasovnikarova et al., 2015) and the ATPase MORC2 to compact chromatin (Douse et al., 2018; Tchasovnikarova et al., 2017). HUSH is a vertebrate-specific chromatin regulator that represses both exogenous and endogenous genetic elements. As well as targeting integrating lentiviruses, HUSH targets full-length transcriptionally-active retrotransposons including LINE-1s (Liu et al., 2018; Robbez-Masson et al., 2018), and cell-type specific genes such as zinc finger transcription factors (*ZNF*s) (Liu et al., 2018; Tchasovnikarova et al., 2015, 2017). HUSH is also recruited, via the DNA-binding protein NP220, to repress expression of unintegrated murine leukemia virus (Zhu et al., 2018). The critical role of HUSH in antiretroviral immunity is highlighted by findings that primate lentiviral accessory proteins Vpr and Vpx target HUSH complex proteins for proteasome-mediated degradation (Chougui et al., 2018; Greenwood et al., 2019; Yurkovetskiy et al., 2018).

In the current model of HUSH-mediated repression, HUSH regulates both reading and writing of H3K9me3 over its targets (Tchasovnikarova et al., 2017). In contrast to classical heterochromatin regulators, some HUSH targets are enriched within transcriptionally active chromatin (Liu et al., 2018; Robbez-Masson et al., 2018). The MPP8 chromodomain binds K9-methylated H3 tail peptide (Kokura et al., 2010) and H3K9-like mimic sequences found in other proteins including ATF7IP, the nuclear chaperone of SETDB1 (Timms et al., 2016). It follows that MPP8 binding to methylated ATF7IP recruits SETDB1 to spread H3K9me3 over HUSH targets (Tsusaka et al., 2018). However, a read-write mechanism for H3K9me3 spreading involving MPP8 and ATF7IP/SETDB1 must be too simplistic, as TASOR and Periphilin are essential to maintain HUSH-dependent lentiviral reporter repression, whereas the MPP8 chromodomain is not (Tchasovnikarova et al., 2015). Hence, a key unanswered question is how TASOR and Periphilin contribute to HUSH repression.

Here we report multiscale biochemical and functional analyses of TASOR that provide new mechanistic insights into how TASOR contributes to epigenetic repression of its targets. We report on the role of Periphilin elsewhere (Prigozhin et al., 2019). TASOR is a 1670-amino acid nuclear protein lacking functional annotations. At the molecular level TASOR remains poorly characterized, apart from its identification as an mRNA binding protein (Castello et al., 2012; Queiroz et al., 2019). Here we show that TASOR is the central assembly platform of HUSH, providing binding sites for MPP8 and Periphilin. Targeted epigenomic profiling experiments support the model that TASOR binds and regulates H3K9me3 specifically over LINE-1 repeats and repetitive exons of transcribed genes. Analysis of TASOR’s domain organization reveals striking homology with the yeast RNA induced transcriptional silencing (RITS) complex subunit Chp1, and in a proteomic screen we find TASOR associates with RNA processing components. Together with observations that transgene transcription enhances HUSH binding (Liu et al., 2018), these data suggest that an RNA intermediate may be important for HUSH activity. Our cellular assays map the specific subdomains of TASOR and MPP8 necessary for HUSH assembly and transgene repression. Structural and biochemical studies reveal that TASOR contains a catalytically inactive poly-ADP ribose polymerase (PARP) domain that is dispensable for assembly and chromatin targeting but critical for epigenetic regulation of target elements. We find that this activity relies on an extended, dynamic loop that is unique in the PARP family. Our data demonstrate TASOR is a pseudo-PARP that governs both HUSH assembly and H3K9me3 deposition over its genomic targets.

## Results

### TASOR binds and regulates H3K9me3 over L1P elements and repetitive exons

We originally identified HUSH as a repressor of lentiviral transgenes (Tchasovnikarova et al., 2015, 2017). Chromatin immunoprecipitation sequencing (ChIP-seq) showed that HUSH also targets hundreds of endogenous genes and transposable elements (Liu et al., 2018; Robbez-Masson et al., 2018; Tchasovnikarova et al., 2017). Notably, transcription promotes target binding by the HUSH subunit MPP8 (Liu et al., 2018), and HUSH loci were found in transcriptionally-active euchromatic regions as defined by epigenetic marks (Liu et al., 2018; Robbez-Masson et al., 2018) and sensitivity to sonication (Becker et al., 2017). These results highlight that H3K9me3 is not restricted to heterochromatin and suggest that sonication of cross-linked chromatin in ChIP protocols could influence analysis of HUSH regulation. TASOR ChIP-seq also displayed low sensitivity (Liu et al., 2018). For these reasons we applied orthogonal and targeted strategies for TASOR epigenomic profiling, CUT&RUN (Skene et al., 2018) and CUT&Tag (Kaya-Okur et al., 2019).

CUT&RUN profiling of H3K9me3 in the presence and absence of TASOR identified 393 TASOR-regulated loci with high resolution and sensitivity (**Fig. 1A, S1A**). The proportion of global H3K9me3 regulated by HUSH – 1% – was comparable to that determined by ChIP-seq (Liu et al., 2018; Tchasovnikarova et al., 2015). We observed a strong association with LINE-1 elements (L1s) in our analysis. Of all the transposable element classes overlapped by TASOR-regulated sites, 86.1% corresponded to L1s of which the majority were primate-specific L1Ps (**Fig. S1B**). TASOR CUT&RUN was unsuccessful, perhaps due to TASOR’s size and poor solubility, but CUT&Tag showed improved sensitivity over ChIP-seq (**Fig. 1A, S1C**). We found that the strongest TASOR CUT&Tag peaks were co-occupied by H3K9me3, consistent with the model that TASOR binds chromatin via MPP8 (**Fig. 1A, S1D**). Secondary H3K9me3-independent binding was also observed at a handful of sites by CUT&Tag (**Fig. S1E**). The resolution afforded by targeted methods enabled boundaries of TASOR binding and associated H3K9me3 deposition to be mapped with precision, revealing that this often coincides with boundaries of L1P sequences (**Fig. 1A, S1D**).

**Fig. 1.**
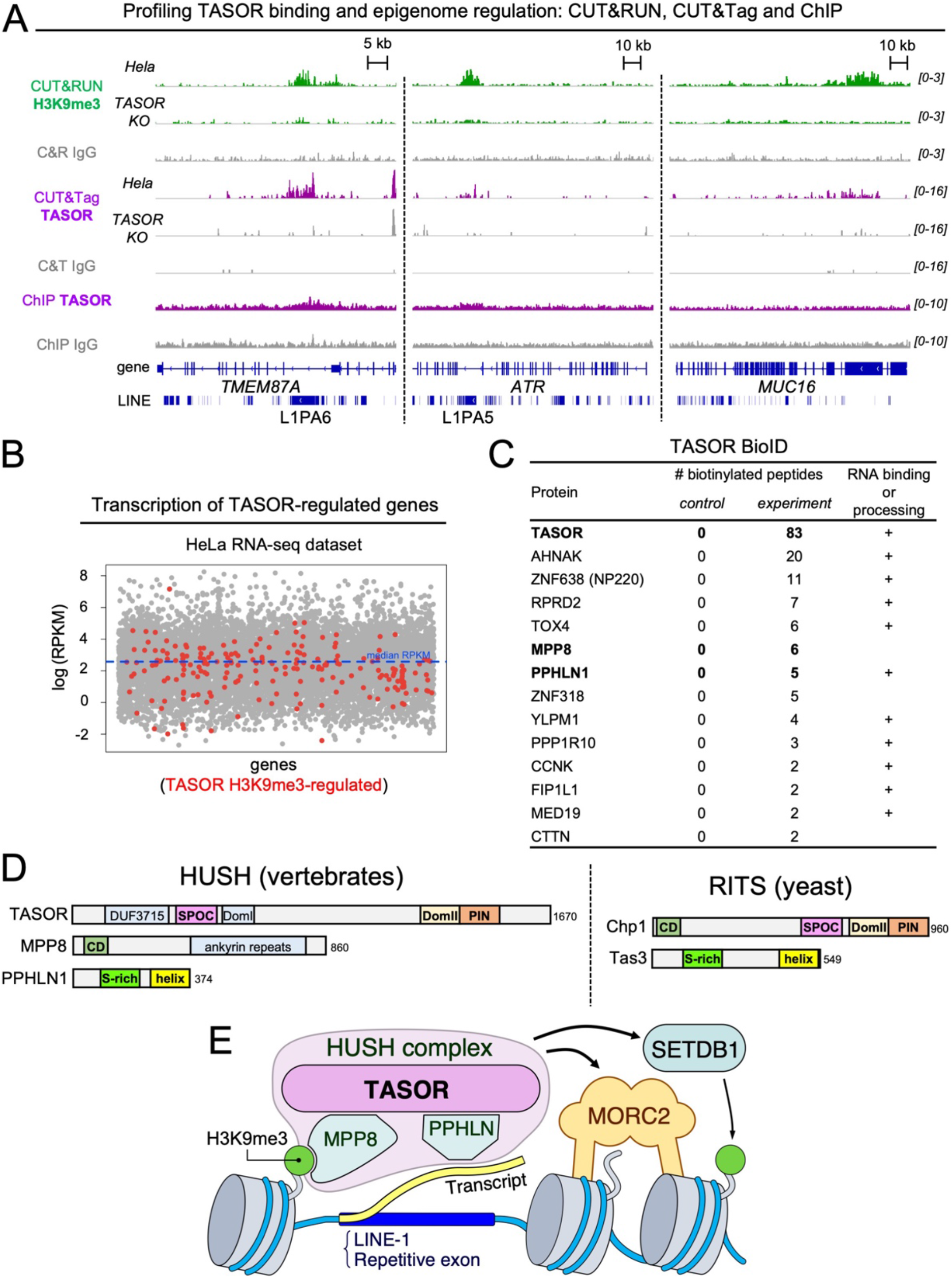
HUSH activities and domain structures resembles those of the yeast RITS complex. (**A**) Genome browser snapshots of TASOR binding and regulation of H3K9me3 in HeLa cells. Shown are CUT&RUN H3K9me3 tracks in TASOR-positive and TASOR-negative cells (green); an IgG negative control track (grey); CUT&Tag and ChIP TASOR tracks in TASOR-positive (purple) and TASOR-negative cells; and an IgG control for each technique (grey). (**B**) Coding genes overlapping with TASOR-regulated H3K9me3 peaks were plotted (red dots) on a scatterplot showing gene transcript levels. Raw data was processed from triplicate RNA-seq data on TASOR-positive HeLa cells, mapped to hg38 (Tchasovnikarova et al., 2017). RPKM, Reads Per Kilobase per Million mapped reads. (**C**) Table of TASOR bioID hits. The HUSH complex is marked in bold. Control refers to KO cells treated with biotin but without expression of BirA-tagged TASOR. (**D**) Predicted domain architecture of *Homo sapiens* HUSH subunits, showing similarity with *Schizosaccharomyces pombe* Chp1 and Tas3. DUF, domain of unknown function. SPOC, Spen ortholog C-terminal. PIN, PilT N-terminus. CD, chromodomain. HUSH, Human Silencing Hub. RITS, RNA-induced transcriptional silencing complex. (**E**) Schematic model of transgene repression by HUSH.

Intronic L1Ps account for most, but not all TASOR-regulated H3K9me3 peaks that overlap with genes. ZNF (zinc finger) genes are significantly enriched (P < 2.2 × 10^−16^, Fisher exact test), consistent with other datasets (Liu et al., 2018; Tchasovnikarova et al., 2015). TASOR-regulated H3K9me3 on *ZNF*s predominantly covers their 3’ exons, which encode repetitive and rapidly-evolving zinc finger arrays (Blahnik et al., 2011; Imbeault et al., 2017). Inspection of other non-L1 targets suggested that TASOR-dependent H3K9me3 deposition may be found over repetitive exons irrespective of gene organization (e.g. *MUC16* exon 3, *BRCA2* exon 11, *C2orf16* exon 1) (**Fig. 1A, S1F**). Repetitive and/or rapidly-evolving genes are potential sources of recombination and genome damage (Aguilera and Gaillard, 2014; Blahnik et al., 2011), a potential shared by L1Ps. Together, these observations suggest that HUSH-dependent H3K9me3 deposition and chromatin compaction could have a genome protective function alongside its established repressive function.

### TASOR binds transcribed genes and associates with RNA processing machinery

Retrovirus and LINE-1 HUSH targets pose a threat to the genome by replication through an RNA intermediate. Induced LINE-1 transcription promoted MPP8 genome binding in K562 cells (Liu et al., 2018) and a subset of HUSH-bound genes were found to show tissue-specific expression (Liu et al., 2018; Robbez-Masson et al., 2018). According to reanalysis of RNA-seq data (Tchasovnikarova et al., 2017), the median expression of genes overlapping TASOR-regulated sites (RPKM = 7.52, n = 228) is comparable to that of all others (RPKM = 7.71, n = 14,211) (**Fig. 1B**). Closer examination supports an association between transcription and H3K9me3 deposition through HUSH. For example, *MUC16* is expressed in HeLa but not K562 cells (Liu et al., 2018) and is only H3K9me3-marked in a TASOR-dependent manner in HeLa cells. *BRCA2* is expressed in HeLa and K562 and is HUSH-modified in both lines (**Fig. 1A**, **S1E**) (Liu et al., 2018). Our data support a model in which transcription of LINE-1s or other repetitive exons correlates with TASOR binding and H3K9me3 deposition over the element.

To investigate potential transient associations made by TASOR on chromatin, we performed proximity-dependent labeling (Bio-ID) using BirA-tagged TASOR in TASOR knockout (KO) cells (**Fig. 1C**). Using this approach we identified peptides from TASOR, MPP8, Periphilin and 10 other chromatin-associated proteins. Among our top hits were matrin-type zinc finger proteins ZNF318 and ZNF638 (NP220). Although NP220 is known to recruit HUSH to unintegrated murine retroviral DNA (Zhu et al., 2018), our data predict additional roles for this interaction in the absence of infection. We also detected RPRD2, a regulator of RNA Polymerase II previously identified alongside TASOR as a repressor of LINE-1s (Liu et al., 2018) and HIV (Liu et al., 2011). 11 of the 14 proteins we identified are annotated RNA binding proteins. Of these, CCNK, MED19, RPRD2 and FIP1L1 have been associated with regulating mRNA processing and RNA polymerase II activity. Association with RNA processing machinery is consistent with observations that HUSH binds and regulates transcriptionally active genomic targets, together with TASOR’s annotation as an mRNA-interacting protein (Castello et al., 2012; Queiroz et al., 2019).

### TASOR is a multidomain protein resembling Chp1 of the yeast RITS complex

Despite its important roles in antiviral defense and vertebrate development, TASOR lacks functional annotations in its 1670-residue sequence apart from a ‘domain of unknown function’ (DUF3715, residues 106-332). Disorder (Jones and Cozzetto, 2015) and structural homology (Kelley et al., 2015; Yang and Zhang, 2015) prediction on the primary sequence of TASOR identified four additional putative domains (**Fig. 1D, S2A,B**). A Spen paralog and ortholog C-terminal (SPOC) beta barrel domain (residues 350-505) is predicted to lie adjacent to DUF3715, while no structural homology was identified for the third ordered region (referred to as DomI, residues 525-633). Residues 1233-1466 exhibit homology to the DomII and PIN domains of the *S. pombe* protein Chp1 (**Fig S2A**). Intriguingly, Chp1 also contains a SPOC domain (**Fig. S2B**) and, like MPP8, an H3K9me3-binding chromodomain. Furthermore, the Chp1 binding partner Tas3 is a small protein that resembles Periphilin (**Fig. 1D**).

Together Chp1 and Tas3 form the core structure of the yeast RNA-induced transcriptional silencing (RITS) complex (Schalch et al., 2011). The striking resemblance of RITS (Chp1-Tas3) domain organization to that of HUSH (**Fig. 1D**) is particularly notable given the functional relationships between the two complexes. In yeast, Tas3 self-associates through C-terminal helical repeats to spread heterochromatic gene silencing (Li et al., 2009a). Periphilin self-association – through disordered and C-terminal helical regions – is likewise required for HUSH repression in human cells (Prigozhin et al., 2019). RITS targets repetitive elements such as centromeres and telomeres, and – as predicted for HUSH (**Fig. 1E**) – mediates deposition of repressive H3K9me3 over repeats in response to transcription (Schalch et al., 2011; Verdel et al., 2004).

### The N-terminal domains of TASOR and C-terminal region of MPP8 are required for HUSH-mediated transgene repression

Next we assessed which TASOR and MPP8 domains are required for HUSH transgene repression. We first generated a panel of TASOR truncation mutants (**Fig. S3A**) and performed genetic complementation assays in TASOR knockout cells harboring a de-repressed GFP lentiviral transgene (**Fig. 2A**). Upon expression of full-length TASOR, HUSH function was restored and the reporter was repressed. However, TASOR deletion mutants lacking the DUF3715, SPOC, or DomI domains were non-functional. A variant lacking the DomII/PIN domains (deletion of residues 1233-1670) complemented the knockout, indicating that this C-terminal region is not required for transgene repression under the conditions tested. We note that DomII/PIN is nonetheless highly conserved in TASOR orthologues, and these domains may therefore play functional roles not captured by our assay. Indeed, the PIN domain of Chp1 was required for repression by RITS at subtelomeric but not centromeric repeats (Schalch et al., 2011). Finally, a variant with additional C-terminal truncation – TASOR(1-1085) – retained function, but TASOR(1-1000) did not.

**Fig. 2.**
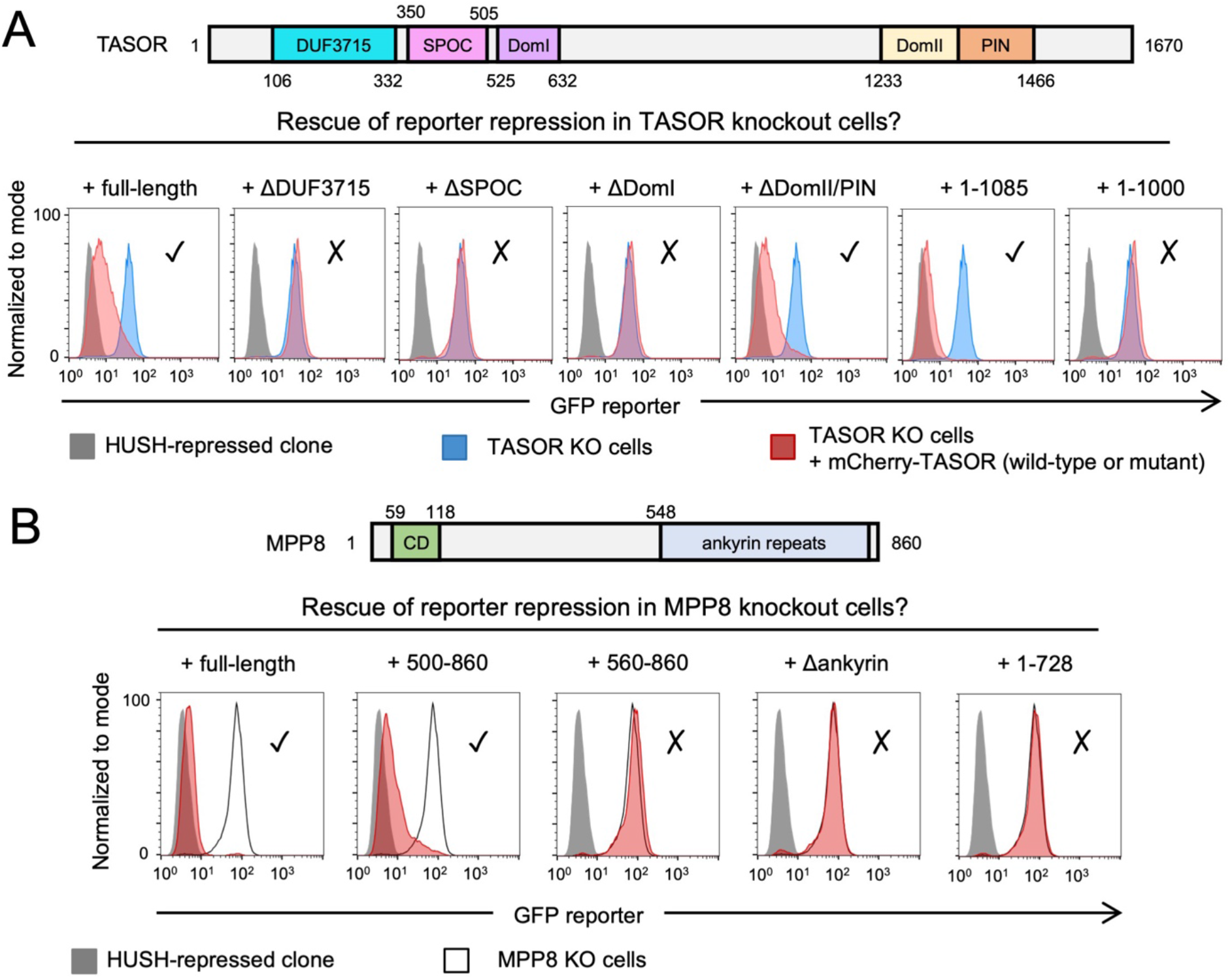
Functional requirements of TASOR and MPP8 domains in HUSH transgene repression. (**A**) Genetic complementation of TASOR knockout cells. Exogenous expression of full-length TASOR or TASOR lacking the DomII/PIN domain restores GFP transgene repression as measured by flow cytometry, but mutants lacking the DUF3715, SPOC or DomI domains are non-functional. TASOR(1-1085) is functional and TASOR(1-1000) is non-functional. (**B)** Genetic complementation assays in MPP8 knockout cells demonstrate that the C-terminal ankyrin repeats are required for function, but MPP8(500-860) is sufficient to complement the knockout. Ticks and crosses summarize functionality in this assay.

The domain structure of MPP8 is comparatively simple: an N-terminal chromodomain separated from C-terminal ankyrin (helix-loop-helix) repeats by a linker. The chromodomain is not strictly essential for the maintenance of HUSH-mediated repression, though a mutation that inhibits H3K9me3 binding delays reestablishment of reporter repression (Tchasovnikarova et al., 2015). Extending this analysis of MPP8, we found that the first 499 amino acids could be removed altogether without further impairing HUSH function. However, truncation to residue 560 caused loss of function, and the C-terminal ankyrin repeats were functionally-required (**Fig. 2B, S3B**).

Together we conclude that the DUF3715, SPOC and DomI domains of TASOR, along with its central linker, are required for transgene repression by HUSH, but the C-terminal 585 residues are dispensable. The C-terminal portion (500-860) of MPP8, which contains the predicted ankyrin repeats, is likewise required.

### TASOR lies at the heart of the HUSH complex

TASOR domains essential for HUSH function could (i) mediate HUSH complex formation through interactions with MPP8 and Periphilin or (ii) have biological activities necessary for repression (or recruit effector proteins with these activities). We first considered the overall role of TASOR in HUSH assembly. Starting from a cell line lacking all three HUSH subunits (Tchasovnikarova et al., 2015), we re-expressed subunits in pairwise combinations and examined their interactions through reciprocal co-immunoprecipitation (co-IP) **(Fig. 3A)**. MPP8 and Periphilin precipitated with TASOR but no binding was detected between MPP8 and Periphilin without TASOR. We conclude that TASOR lies at the heart of the core HUSH complex.

**Fig. 3.**
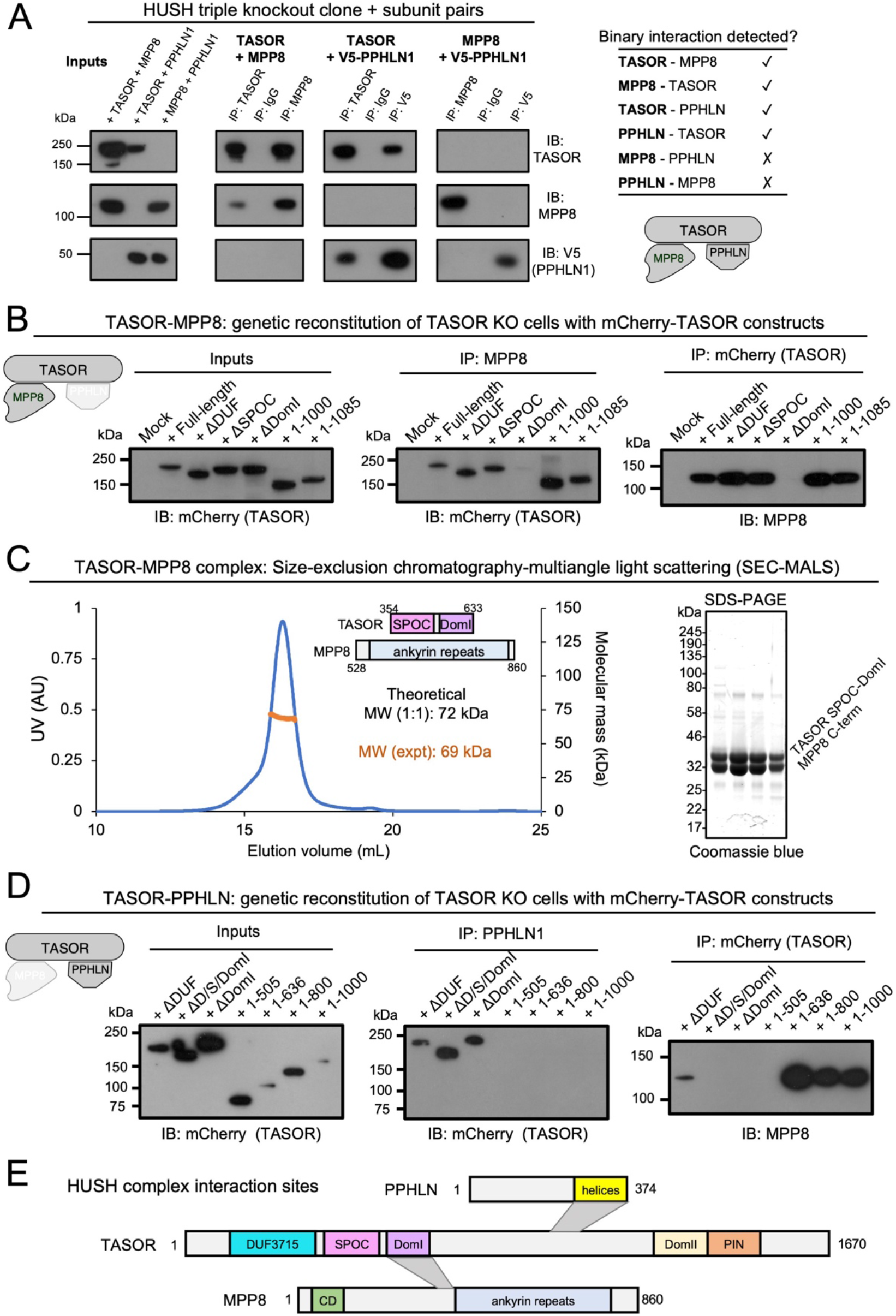
TASOR lies at the heart of the HUSH complex. (**A**) TASOR interacts separately with MPP8 and Periphilin to mediate assembly of the HUSH complex. Pairwise combinations of HUSH subunits were exogenously expressed in HUSH triple KO cells, and interactions detected by co-immunoprecipitation (co-IP) followed by immunoblot. (**B**) TASOR DomI mediates the interaction with MPP8. The indicated TASOR truncation mutants were exogenously expressed in TASOR KO cells and their association with endogenous MPP8 assessed by co-IP. (**C**) SEC-MALS of recombinant TASOR(354-633)-MPP8(528-860) complex (theoretical MW, 72 kDa for a 1:1 complex) with representative SDS-PAGE gel of SEC peak fractions (inset). (**D**) The central linker of TASOR mediates the interaction with Periphilin. (**E**) Summary of core HUSH interaction sites.

To map binding sites, we performed co-IP of endogenous MPP8 with tagged TASOR truncation variants or vice versa. TASOR DomI was required for MPP8 binding regardless of which protein was immunoprecipitated **(Fig. 3B)**. The adjoining SPOC domain was dispensable for MPP8 binding in cells, which was unexpected since SPOC domains from other transcriptional regulators like yeast Chp1 form stabilizing protein-protein interactions through an exposed hydrophobic patch (Ariyoshi and Schwabe, 2003; Schalch et al., 2011). Homology modelling suggests that this hydrophobic patch is also present in TASOR. Attempts to isolate recombinant TASOR SPOC or DomI domains, or TASOR-MPP8 complexes based on DomI only, resulted in poor yields of soluble protein. However, we were able to purify a stable, 1:1 TASOR-MPP8 complex following recombinant co-expression of a tandem TASOR SPOC-DomI construct (residues 354-633) and the MPP8 C-terminus (residues 528-860) (**Fig. 3C**). Together these data support a binding mode in which the TASOR SPOC domain stabilizes DomI and that the interaction with MPP8 occurs through a largely hydrophobic interface. Since MPP8(500-860) is functional but MPP8(560-860) is non-functional, our data suggest that the minimal TASOR binding site lies between MPP8 residues 500-560.

To assess the TASOR-Periphilin interaction we immunoprecipitated endogenous Periphilin and blotted for mCherry-tagged TASOR constructs. Periphilin pulled down TASOR mutants lacking the DUF3715, SPOC and DomI domains, but not TASOR(1-1000) or TASOR variants with longer C-terminal truncations (**Fig. 3D**). Given that TASOR(1-1085) is functional, these data suggest that the Periphilin binding site lies in residues 1000-1085. Indeed we report elsewhere the crystal structure of a minimal TASOR-Periphilin complex, showing that TASOR(1014-1095) contributes directly to binding the Periphilin C-terminus (Prigozhin et al., 2019). Together these results delineate the biochemical requirements for HUSH assembly, and suggest that the central portion of TASOR, spanning residues 350-1085, is a sufficient assembly scaffold (**Fig. 3E**).

### TASOR DUF3715 is a PARP domain

Having established that TASOR acts as the core member of HUSH and defined the domains necessary for assembly, TASOR’s N-terminal DUF3715 stood out as significant for several reasons: (i) it is required for transgene repression in a manner independent of assembly; (ii) it is the domain that differentiates TASOR from *S. pombe* Chp1 and (iii) it contains the embryonic lethal mouse mutation (L130P) – underlining its key functional role at an organismal level. Furthermore, bioinformatic analysis suggested that DUF3715 resembles a poly ADP-ribose polymerase (PARP) catalytic domain. Functions of the PARP family, which contains at least 18 members in humans, include response to genome damage and viral infection (Gupte et al., 2017). This is notable given TASOR’s role as a LINE-1 and viral repressor.

We therefore aimed to study the structure and biochemical properties of DUF3715. Despite extensive trials, we could not induce the wild-type DUF3715 to crystallize. We used NMR spectroscopy to gain insight into its structural and dynamic properties and recorded well-resolved ^1^H, ^15^N correlation spectra on ^15^N-labelled protein (**Fig. 4A**). To obtain peak assignments we required sidechain deuteration and expressed the domain in media prepared with deuterated water (^2^H_2_O). Upon purification in (^1^H) aqueous solvents, several peaks in the spectrum were missing, consistent with a rigid core in which the rate of amide H-D exchange is exceptionally slow (>5 days). A partial denaturation-refolding protocol enforced exchange of core amide deuterons (**Fig. S4A**), enabling assignment of 184 (of 221) non-proline backbone amide resonances. Secondary structure prediction from chemical shifts (Shen et al., 2009) confirmed an α/β PARP fold with loop insertions (**Fig. S4B**).

**Fig. 4.**
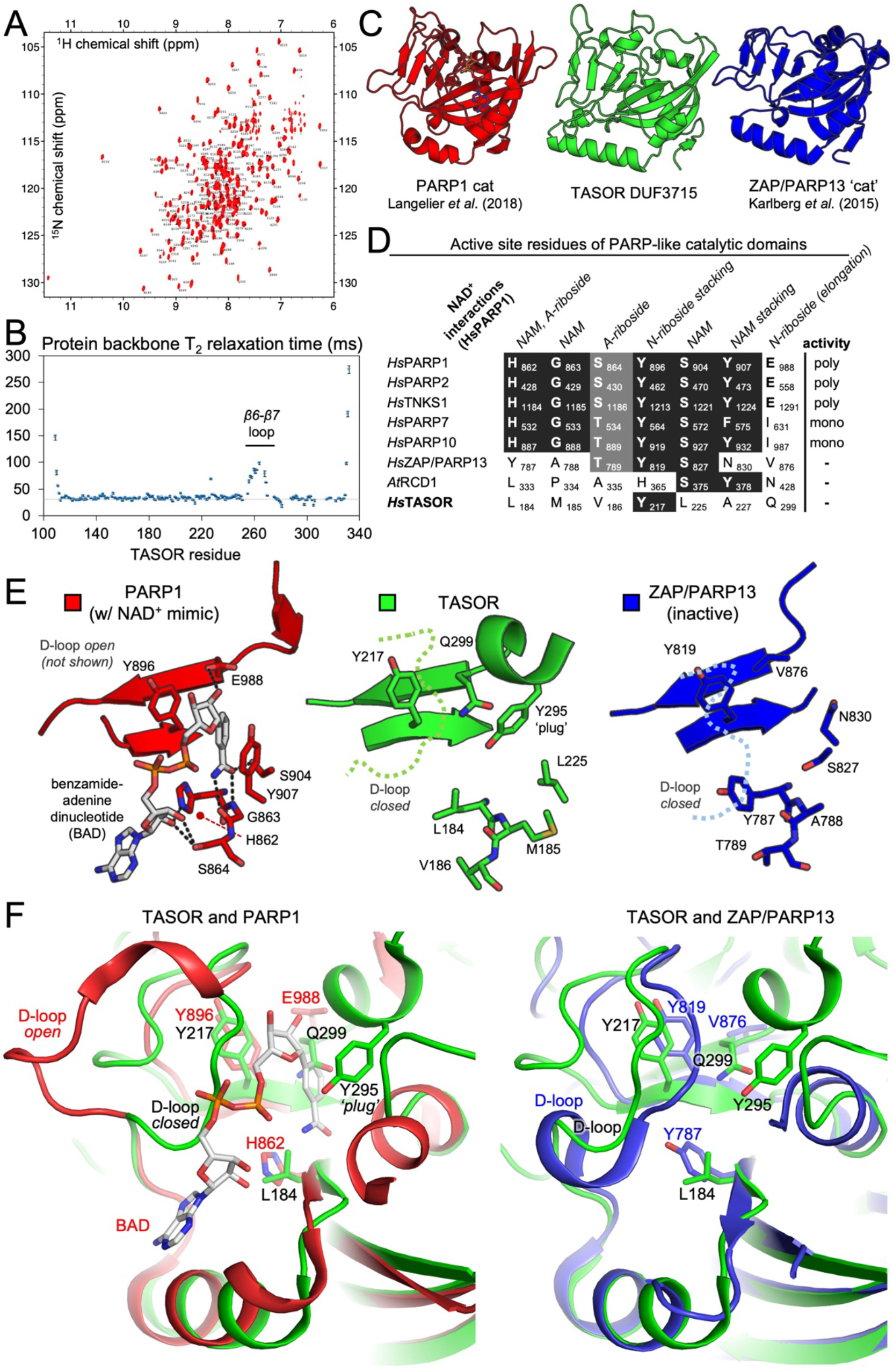
TASOR DUF3715 is a PARP domain. (**A**) Assigned ^1^H-^15^N BEST TROSY spectrum of ^15^N-labelled TASOR(106-332) at 293 K. (**B**) Backbone T_2_ relaxation times of TASOR(106-332). The disordered β6-β7 loop that was removed to promote crystallization is marked. (**C**) Overall structures of PARP domains of human PARP1 (red), TASOR (green) and ZAP (blue). (**D**) Alignment of PARP active sites and NAD+ binding residues for a selection of PARP family members. NAM, nicotinamide; A-riboside, adenosyl-riboside; N-riboside, nicotinamide-riboside. (**E**) and (**F**) Comparison of active sites of the structures shown in panel C, with key features marked.

### Overall structure and dynamics of TASOR PARP domain

NMR assignments allowed us to measure T_1_ and T_2_ relaxation times to investigate domain dynamics (**Fig. 4B**). The average T_1_/T_2_ ratio is proportional to overall tumbling correlation time τ_C_, which we determined to be ~15 ns. This value suggests DUF3715 is largely monomeric in solution at 250 μM, though the oligomeric status of full-length TASOR remains unknown. Elevated T_2_ relaxation times for residues 255-270 (the loop connecting strands β6-β7) were indicative of motions faster than overall tumbling (τ_C_), in the ns-ps timescale (Kay et al., 1989) – commonly described as disorder (**Fig. 4B**). A TASOR variant with part of this disordered loop deleted (Δ261-269) retained transgene repression activity **(Fig. S4C)**.

Hypothesizing that β6-β7 loop disorder precluded crystallization of the wild-type domain, we purified a variant with the Δ261-269 deletion. Validating our design, crystals were readily grown of native and SeMet-labelled samples (**Fig. S4D**), enabling X-ray structure determination of TASOR DUF3715 to a resolution of 2.0 Å by single-wavelength anomalous dispersion (SAD) phasing (**Table S1**). Comparison with published structures using the Dali server (Holm and Laakso, 2016) confirmed similarity of the overall fold with PARPs from various subclasses (**Fig. 4C**) including canonical poly-(*Hs*PARP1, PDB:6BHV), mono-(*Hs*PARP10, PDB:3HKV) and catalytically-inactive (*Hs*ZAP/PARP13, PDB:2X5Y) ADP-ribosyl transferases, and the inactive plant PARP homologue RCD1 (*At*RCD1, PDB:5NGO).

### The TASOR PARP domain lacks an NAD+ binding site

PARPs use NAD^+^ as a cofactor to catalyze addition of one or more ADP-ribosyl units onto target proteins, although some PARPs have lost this activity. Since mechanistic studies of ADP-ribosylation are hampered by a lack of understanding of PARP substrates, crystal structures have been a useful means to classify the family into active and inactive members (Wahlberg et al., 2012). The structure of PARP1 catalytic domain in complex with non-hydrolysable NAD^+^ analogue benzamide adenosine dinucleotide (BAD) provides a near-complete picture of interactions made during NAD^+^ binding (Langelier et al., 2018). We tabulated amino acids making ligand contacts in the PARP1-BAD structure, including the canonical histidine-tyrosine-glutamate triad required for poly-ADP ribosylation (**Fig. 4D, E**). Substitution of the glutamate, as in PARP7 and PARP10, removes capacity for chain elongation, thus limiting these enzymes to mono-ADP-ribosylation (Kleine et al., 2008). The other residues involved in NAD^+^ binding are conserved in active PARPs, regardless of whether activity is mono- or poly-ADP ribosylation. By contrast, the human zinc finger antiviral protein (*Hs*ZAP, or PARP13) and *Arabidopsis* RCD1 lack two or more of the NAD^+^-binding residues and are catalytically inactive (Karlberg et al., 2015; Wirthmueller et al., 2018). In TASOR the key residues required for catalysis are even more degenerated than in ZAP or RCD1: all but one of them are replaced by hydrophobic amino acids (**Fig. 4D-F**). TASOR shares a further similarity with ZAP/PARP13, in that the equivalent of its active site loop or D-loop (TASOR residues 200-210) adopts a closed conformation relative to PARP1, in which it is open (Karlberg et al., 2015; Langelier et al., 2018) (**Fig. 4E,F**). Together with a short helix spanning residues 294-298, these features occlude the TASOR ligand-binding pocket and the sidechain of Tyr295 plugs the nicotinamide binding cleft. Based on our structure, TASOR therefore lacks the chemical functionality and physical space to bind the NAD^+^ cofactor.

### TASOR is a pseudo-PARP that binds weakly to ssRNA

Consistent with our structural data illustrating a degenerate active site, point mutations removing the only remaining conserved amino acid (Y217A) or restoring a key NAD^+^ binding residue (L184H) did not affect TASOR-dependent transgene repression in cells (**Fig. 5A,B**). Differential scanning fluorimetry (DSF) experiments showed that while the domain was stable in isolation (T_m_ 51.5°C), no change in T_m_ was observed upon addition of up to 1.5 mM benzamide, a promiscuous NAD^+^-mimic and PARP inhibitor (**Fig. 5C**). By contrast, the catalytic domain of PARP1 showed robust concentration-dependent stabilization as expected (Langelier et al., 2018). We then performed a gel-based poly-ADP-ribosylation (PARylation) assay by mixing recombinant full-length human PARP-1 with the TASOR PARP domain or negative control BSA. While PARP-1 robustly auto-PARylated in the presence of NAD^+^ and dsDNA, shown by a high molecular weight smear in the Coomassie-stained gel, we did not detect significant evidence of TASOR modification or auto-modification under the conditions tested (**Fig. 5D**). We conclude that TASOR’s active site is non-catalytic in isolation, as in human ZAP/PARP13 (Karlberg et al., 2015).

**Fig. 5.**
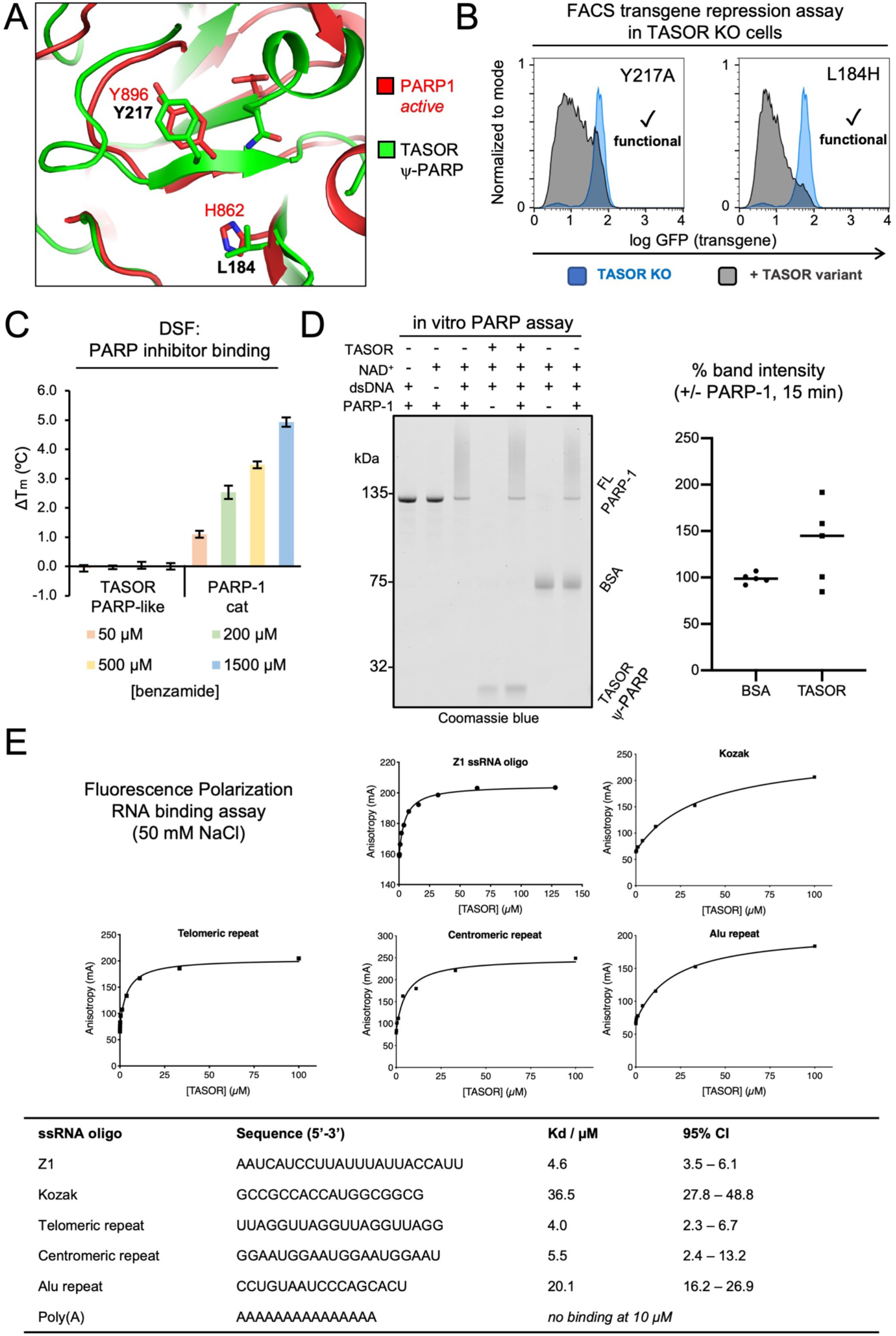
TASOR is a pseudo-PARP. (**A**) Active site comparison of PARP1 and TASOR, showing key functional residues. (**B**) Transgene repression assays show that mutation of the degenerated TASOR PARP active site does not affect TASOR function. (**C**) TASOR PARP domain does not bind promiscuous PARP inhibitor benzamide. DSF of three replicates. (**D**) Full-length PARP1 auto-PARylates in the presence of dsDNA and NAD+ but TASOR PARP domain is not significantly affected. (**E**) Binding of 5’(6-FAM)-labelled ssRNA oligonucleotides to TASOR(106-332) by fluorescence polarization (FP), with a table summarizing the binding data.

ZAP – the other non-catalytic human PARP – functions by binding and degrading target viral RNAs (Guo et al., 2007). Because of this and the association between TASOR and transcription (**Fig. 1**) we were interested to note during purification that recombinant TASOR(106-332) coeluted with expression host RNA. Bound RNA could be removed by treatment with high salt and RNase, but not DNase (**Fig. S4E**). The purified domain bound to various short ssRNA ligands with micromolar affinity at 50 mM NaCl, with limited sequence specificity (**Fig. 5E**). This weak interaction was screened at higher salt concentrations, suggesting that binding is driven primarily by electrostatics. Nonetheless it is plausible that the PARP domain contributes to RNA binding by full-length TASOR, or that the TASOR PARP domain binds an as yet unidentified RNA sequence with high affinity. RNA binding is consistent with the annotation of TASOR (along with Periphilin) as part of the HeLa mRNA interactome (Castello et al., 2012; Queiroz et al., 2019), and association with mRNA processing machinery (**Fig. 1C**). It may be that as in ZAP, other domains in HUSH confer greater affinity or specificity to mRNA binding. Taken together, our structural and biochemical data show that TASOR contains a catalytically inactive PARP domain and that, like ZAP and RCD1, TASOR may be considered a pseudo-PARP (Ribeiro et al., 2019).

### Pseudo-PARP loop β8-β9 undergoes concerted motions and is required for HUSH activity

Topological differences between our structure and the canonical PARP fold are most pronounced in loops. In particular the loop connecting the final two strands (β8 and β9 in TASOR) spans 14 residues in TASOR (Tyr303-His316), compared to 5 residues in all annotated PARP family members. In TASOR the extended β8-β9 loop has elevated *B*-factors in the crystal structure, and broad NMR peaks caused by fast T_2_ relaxation (**Fig. 6A**). The latter is suggestive of conformational changes occurring on timescales greater than τ_C_ (i.e. ms-μs). Such concerted motions require a higher activation energy and often correlate with functionally-relevant processes like conformational exchange (Kay et al., 1989). This appears to hold true for TASOR: variants Δ307-312 and Y305A were non-functional in our transgene repression assay (**Fig. 6B**).

**Fig. 6.**
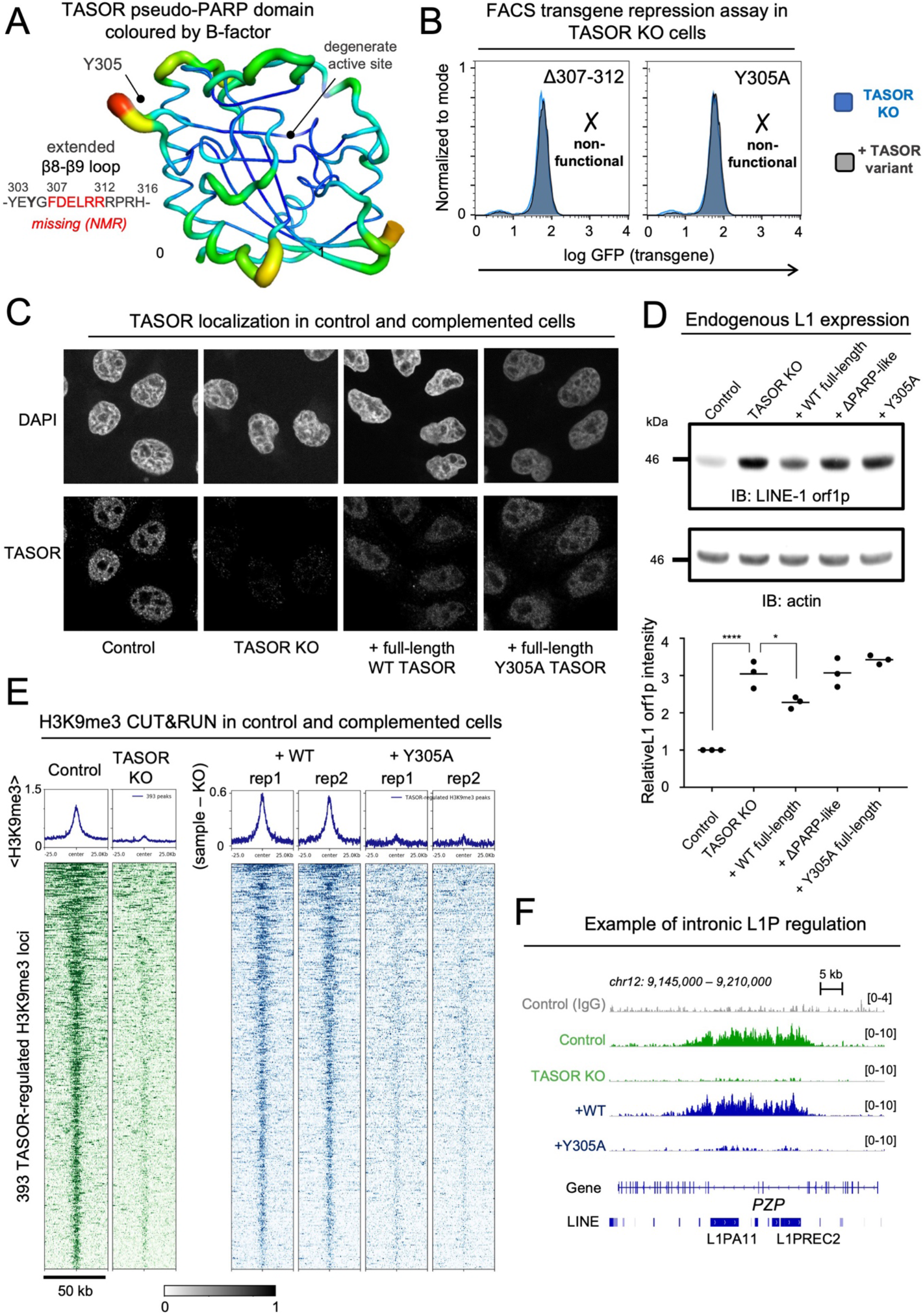
Y305A in the TASOR pseudo-PARP domain abrogates HUSH transgene repression, LINE-1 restriction and TASOR-dependent H3K9me3 deposition genome-wide. (**A**) Structure of TASOR pseudo-PARP domain colored by B-factor. The sequence of the β8-β9 loop is highlighted. These residues were broadened by fast T2 relaxation in NMR spectra. (**B**) Δ307-312 or Y305A mutations in the β8-β9 loop cause loss of TASOR GFP transgene repression. (**C**) Confocal immunofluorescence of nuclei of indicated cell lines with DAPI or TASOR staining. Size of each image, 72 × 72 μm. (**D**) TASOR KO causes derepression of LINE-1 orf1 expression. This repression is partly rescued by full-length WT TASOR but not TASOR ΔPARP nor Y305A. *, p < 0.05; ****, p < 10-4 that KO and control or complemented cell lines are the same (n=3). (**E**) Heatmaps showing summary of H3K9me3 complementation CUT&RUN experiment. Examples of H3K9me3 profiles in control and TASOR KO cells at 393 TASOR-regulated loci. Y305A mutation causes loss of H3K9me3 (sample – KO) at 393 TASOR-regulated loci. Both replicates are shown. (**F**) Genome browser snapshot of CUT&RUN data showing rescue of H3K9me3 regulation over two intronic L1P elements in gene *PZP* for WT but not Y305A TASOR.

We then aimed to further dissect why the Y305A mutation abrogates transgene repression. First, we purified the Y305A mutant to confirm that it did not cause domain misfolding (**Fig. S5A**). DSF showed that the mutant (T_m_ 48.0°C) was modestly destabilized compared to WT (T_m_ 51.5°C) but fully folded at physiological temperature. In cells we confirmed that the Y305A mutant remained localized in the nucleus (**Fig. 6C**). Moreover, the ΔPARP deletion mutant remained chromatin-associated (**Fig. S5B**) and both TASOR Y305A and ΔPARP also interacted with the same set of cellular proteins as WT in BioID experiments (**Fig S5C**). Together, these data are consistent with a model in which the pseudo-PARP is not required for genome localization or HUSH assembly, although it remains possible that the domain contributes to TASOR targeting under certain conditions. Finally we asked whether the Y305A mutation affected LINE-1 repression and TASOR-regulated H3K9me3 deposition. Expression of LINE-1 orf1p in TASOR KO cells was partially re-repressed by expression of WT TASOR but not the ΔPARP or Y305A variants (**Fig. 6D**). CUT&RUN profiling showed that the Y305A point mutation caused a near-complete loss of H3K9me3 deposition at the 393 sites we defined to represent TASOR-regulated H3K9me3, an effect functionally equivalent to TASOR KO (**Fig. 6E,F**). We therefore conclude that the extended, dynamic β8-β9 loop – conserved in TASOR but unique among the PARP family – is necessary not only for reporter transgene repression but also genome-wide H3K9me3 deposition and LINE-1 restriction by HUSH.

## Discussion

This study provides a molecular characterization of TASOR and how it contributes to HUSH function. We have shown that TASOR is as an assembly platform for MPP8 and Periphilin, and we have identified the minimal molecular determinants of HUSH complex assembly.

TASOR DomI interacts with MPP8(500-560), and TASOR(1014-1085) interacts with Periphilin(285-374) (**Fig. 3**, see also (Prigozhin et al., 2019)).

We report striking homology in the domain organization of the HUSH complex and the yeast RNA-induced transcriptional silencing (RITS) complex (**Fig. 1**). The core subunits of each complex, TASOR and Chp1, both contain a SPOC domain and C-terminal DomII/PIN domains. Both complexes also contain N-terminal chromodomains, in Chp1 and MPP8 respectively, that recognize H3K9me3-marked chromatin. Chp1 binding partner Tas3 also resembles HUSH subunit Periphilin: both are largely unstructured with low-complexity sequences and C-terminal helical repeats. Homology between RITS and HUSH is particularly interesting in light of several functional similarities between the two complexes. Both HUSH and RITS affect H3K9me3 deposition over targets, and rely on self-association (of Tas3 or Periphilin) to spread this epigenetic mark (Li et al., 2009a; Prigozhin et al., 2019). In RITS H3K9me3 deposition is induced by a positive feedback loop based on recognition of nascent repetitive transcripts. Observations that MPP8 genome binding is enhanced by LINE-1 transcription (Liu et al., 2018) and that TASOR and Periphilin are mRNA binding proteins (Castello et al., 2012; Prigozhin et al., 2019; Queiroz et al., 2019) are therefore pertinent. The functional relationship between HUSH and RITS is further supported by our epigenomic profiling of TASOR-regulated H3K9me3, which strengthens the notion that HUSH targets for H3K9 methylation in human cells are primarily intronic L1P repeats, or repetitive exons in transcribed genes.

HUSH target elements have in common the potential to cause genome damage if incorrectly processed. HUSH may therefore play a role in controlling the rate of transcription of these elements. RNA Polymerase II (Pol II) is known to transcribe H3K9me3-marked and repetitive regions more slowly (Saint-André et al., 2011; Veloso et al., 2014). We have reported elsewhere that Periphilin binds RNA and aggregates through a low-complexity sequence (Prigozhin et al., 2019). The formation of Periphilin-RNA aggregates could potentially physically impede transcriptional elongation by Pol II. Our protein-protein interaction screen identified several mRNA processing factors as TASOR binders (**Fig. 1**). One notable example is RPRD2, another LINE-1 repressor (Liu et al., 2018), which regulates transcription by directly binding Pol II (Ni et al., 2014). HUSH-dependent H3K9me3 could similarly reduce transcription rates to prevent R-loop formation at genomic regions prone to instability (Aguilera and Gaillard, 2014), or to ensure correct mRNA processing (e.g. splicing). Indeed, intronic LINE-1s are thought to function as hubs of transcriptional repression that protect long mammalian introns from improper splicing (Attig et al., 2018). Local chromatin compaction by HUSH cofactor MORC2 may provide an additional protective barrier of repression (Tchasovnikarova et al., 2017). Loss of these repressive barriers should cause transcription rates to increase, potentially leading to improper processing and genome instability. This may explain the developmental arrest observed in mouse TASOR mutants at the onset of gastrulation.

A potential role for HUSH in genome protection is consonant with our discovery of a pseudo-PARP domain in TASOR. PARPs are central regulators of genome stability, which can be compromised by inappropriate recombination or retrotransposition. We found that the TASOR pseudo-PARP is critical for transgene repression (**Fig. 2**), despite being dispensable for HUSH assembly. Unlike canonical PARPs, TASOR appears from our crystal structure to be incompetent for catalysis (**Fig. 4,5**). There are intriguing parallels between TASOR and ZAP (PARP13), which is also catalytically inactive, because ZAP functions by influencing the destruction of target RNAs (Schwerk et al., 2019; Takata et al., 2017) and also inhibits LINE-1 retrotransposition (Moldovan and Moran, 2015). We have identified that TASOR activity relies on an extended loop unique among the PARP family, and a single point mutation (Y305A) in this loop is sufficient to abolish transgene repression, endogenous LINE-1 restriction and genome-wide H3K9me3 deposition (**Fig. 6**). Structural studies with larger fragments of HUSH in complex with chromatin or RNA will be necessary to illuminate the basis for this loss of function. In light of our NMR data we speculate that the Y305A mutation inhibits a conformational change within the chromatin-engaged HUSH complex, leading to a loss of repression and H3K9me3 deposition.

Our findings have implications for how HUSH regulation is established and maintained over newly integrated genetic elements such as retroviruses. This is clinically important in the context of HIV latency and gene therapy (Chougui and Margottin-Goguet, 2019). We reported previously that HUSH represses transgenes that integrate into H3K9me3-marked chromatin. However, the requirement for the pseudo-PARP domain and Periphilin self-association (Prigozhin et al., 2019) in HUSH function (but not assembly) underscores that H3K9me3 reading and writing is insufficient to explain HUSH activities. Whether HUSH has H3K9me3-independent modes of recruitment to target sequences also remains to be resolved. Silencing of unintegrated murine retroviral DNA by HUSH requires NP220 (Zhu et al., 2018), a DNA- and RNA-binding protein thought to be at least partly sequence-specific for cytidine clusters (Inagaki et al., 1996). We find that NP220 and another matrin-type ZNF (ZNF318) interact with TASOR in the absence of virus, raising the possibility that multiple specific adaptors could recruit HUSH in different contexts. We note that although nucleotide sequences are one source of specificity in epigenetic repression, in the case of HUSH, specificity could also arise from RNA structure or the repetitiveness of a nucleotide sequence.

## Methods

### Cell culture

HeLa and HEK293T cells (ECACC) were grown in IMDM or DMEM plus 10% fetal calf serum (FCS) and penicillin/streptomycin (100 U/mL). Cell lines were routinely tested for mycoplasma contamination using the MycoAlert detection kit (Lonza).

### Antibodies

The following primary antibodies were used: rabbit α-TASOR (Atlas HPA006735, RRID:AB_1852384 for ChIP, CUT&Tag and Western blot); rabbit α-TASOR (abcam ab224393, for microscopy); rat α-mCherry (Thermo Fisher Scientific, M11217, RRID:AB_2536611); rabbit α-MPP8 (Proteintech, 16796-1-AP, RRID:AB_2266644); rabbit-α-V5 (Abcam, ab27671, RRID:AB_471093); mouse α-Myc (Abcam, ab32, RRID:AB_303599); mouse α-FLAG (Millipore Sigma, F1804, RRID:AB_262044; rabbit α-PPHLN1 (Abcam, ab69569, RRID:AB_1269877); rabbit α-H3K9me3 (abcam ab8898, RRID:AB_306848, for CUT&RUN and CUT&Tag); rabbit α-H3K27me3 (CST C36B11, RRID:AB_2616029, for CUT&RUN positive control); guinea pig α-rabbit IgG (CSB-PA00150E1Gp, for CUT&Run and CUT&Tag); rabbit IgG (CST 2729, RRID:AB_1031062 for ChIP negative control); rabbit α-ORF1p (CST D3W9O, RRID:AB_2800129); mouse α-β-actin (abcam ab8226, RRID:AB_306371). Dilutions were made according to manufacturer recommendations unless otherwise stated.

### Lentiviral expression

Exogenous expression of TASOR variants and other HUSH components was achieved using the lentiviral expression vectors pHRSIN-pSFFV-GFP-WPRE-pPGK-Hygro, pHRSIN-pSFFV-GFP-WPRE-pPGK-Blasto or pHRSIN-pSFFV-GFP-pPGK-Puro with mCherry-TASOR (full-length or mutant), HA-BirA-TASOR (full-length or mutant), V5-PPHLN1 or myc-SETDB1 cassettes inserted in place of GFP. Lentivirus was generated through the triple transfection of HEK 293T cells with the lentiviral transfer vector plus the two packaging plasmids pCMVΔR8.91 and pMD.G using TransIT-293 transfection reagent (Mirus) as recommended by the manufacturer. Viral supernatant was typically harvested 48 h post-transfection, cell debris removed using a 0.45 μm filter, and target cells transduced by spin infection at 800 g for 1 h. Transduced cells were selected with hygromycin (100 μg/mL), blasticidin (5 μg/mL) or puromycin (2 μg/mL).

### Co-immunoprecipitation and Western blotting

For co-immunoprecipitation, cells were lysed in 1% NP-40 in TBS plus 10 mM iodoacetamide, 0.5 mM phenylmethylsulfonyl fluoride (PMSF) and benzonase (Sigma-Aldrich) for 30 min. Protein A and IgG-sepharose resin was added to the lysates along with primary antibody. The suspension was incubated for 2 h at 4°C and the resin was washed three times in lysis buffer. For Western blotting, cells were lysed with lysis buffer containing 1% SDS instead of 1% NP-40. For SDS-PAGE analysis, resins or lysates were heated to 70°C in SDS sample buffer for 10 min and run on a polyacrylamide gel. Gels were blotted onto PVDF membranes (Millipore). Blots were blocked in 5% milk in PBS, 0.2% Tween-20 and incubated with primary antibody diluted in blocking solution. As the Periphilin antibody was unable to detect its epitope under NP-40 lysis conditions, we used a mouse antibody against the V5 tag (Abcam, ab27671) as the primary antibody for Periphilin. For TASOR, the primary antibody was rabbit α-TASOR (Atlas, HPA006735). Blots were imaged with West Pico or West Dura (Thermo Fisher Scientific).

### Flow cytometry

Cells were fixed in 1% PFA and analysed on a FACSCalibur or a FACSFortessa instrument (BD). Data was analysed using FlowJo software. For cell sorting, cells were resuspended in PBS + 2% FCS and an Influx cell sorter (BD) was used.

### CUT&RUN

We followed the protocol detailed by the Henikoff lab (Skene et al., 2018). Briefly, 1-2.5 × 10^5^ cells (per antibody/cell line combination) were washed twice (20 mM HEPES pH 7.5, 150 mM NaCl, 0.5 mM spermidine, 1x Roche complete protease inhibitors) and attached to ConA-coated magnetic beads (Bangs Laboratories) that had been pre-activated in binding buffer (20 mM HEPES pH 7.9, 10 mM KCl, 1 mM CaCl_2_, 1 mM MnCl_2_). Cells bound to the beads were resuspended in 50 μL buffer (20 mM HEPES pH 7.5, 0.15 M NaCl, 0.5 mM Spermidine, 1x Roche complete protease inhibitors, 0.02% w/v digitonin, 2 mM EDTA) containing primary antibody (1:100 dilution). Incubation proceeded at 4 °C for at least 2 h (usually overnight) with gentle shaking. Tubes were then placed on a magnet stand to allow removal of unbound antibody, and washed three times with 1 mL digitonin buffer (20 mM HEPES pH 7.5, 150 mM NaCl, 0.5 mM Spermidine, 1x Roche complete protease inhibitors, 0.02% digitonin). After the final wash, pA-MNase (35 ng per tube, a generous gift from Steve Henikoff) was added in a volume of 50 μL of the digitonin buffer and incubated with the bead-bound cells at 4 °C for 1 h. Beads were washed twice, resuspended in 100 μL of digitonin buffer, and chilled to 0-2 °C. Genome cleavage was stimulated by addition of 2 mM CaCl_2_ (final), briefly vortexed and incubated at 0 °C for 30 min. The reaction was quenched by addition of 100 μL 2x stop buffer (0.35 M NaCl, 20 mM EDTA, 4 mM EGTA, 0.02% digitonin, 50 ng/μL glycogen, 50 ng/μL RNase A, 10 fg/μL yeast spike-in DNA (a generous gift from Steve Henikoff)) and vortexing. After 10 min incubation at 37 °C to release genomic fragments, cells and beads were pelleted by centrifugation (16,000 g, 5 min, 4 °C) and fragments from the supernatant purified with a Nucleospin PCR clean-up kit (Macherey-Nagel). Experimental success was evaluated by capillary electrophoresis (Agilent) with this material and the presence of nucleosome ladders for histone modifications (H3K27me3 or H3K9me3) but not for IgG controls. Illumina sequencing libraries were prepared using the Hyperprep kit (KAPA) with unique dual-indexed adapters (KAPA), pooled and sequenced on a HiSeq4000 or NovaSeq6000 instrument. Paired-end reads (2×150) were aligned to the human and yeast genomes (hg38 and R64-1-1 respectively) using Bowtie2 (--local –very-sensitive-local –no-mixed –no-discordant –phred33 -I 10 -X 700) and converted to bam files with samtools (Langmead and Salzberg, 2012; Li et al., 2009b). Conversion to bedgraph format and normalization was done with bedtools genomecov (-bg - scale), where the scale factor was the inverse of the number of reads mapping to the yeast spike-in genome (Quinlan and Hall, 2010). Peaks were called in SEACR (Meers et al., 2019) (stringent, norm options) for experimental samples against IgG controls. The RUVseq package was used to remove unwanted variation prior to differential binding analysis with edgeR (Risso, 2015; Robinson et al., 2009). TASOR-regulated H3K9me3 peaks were defined as those under a cutoff in the MA plot (log_2_ fold-change < −1) of KO vs control cells. CUT&RUN experiments to assess H3K9me3 regulation were done with four replicates (control and TASOR KO cells, used to define TASOR-regulated peaks) or two replicates (WT and Y305A complemented cells, used to assess H3K9me3 complementation over TASOR-regulated peaks). Normalized bigwig files were generated (UCSC), displayed in IGV (Robinson et al., 2011) and heatmaps plotted with deepTools computeMatrix and plotHeatmap commands (Ramírez et al., 2014).

### ChIP-seq

Cells (10 million per IP) were washed once in PBS, resuspended in growth medium, and then cross-linked in 1% formaldehyde for 10 min. The reaction was quenched by adding glycine to a final concentration of 0.125 M for 5 min before the cells were lysed in cell lysis solution (10 mM HEPES pH 7.5, 85 mM KCl, 0.5% IGEPAL). Nuclei were pelleted by centrifugation, and then resuspended in nuclear lysis solution (50 mM Tris pH 8.1, 10 mM EDTA, 1% SDS) for 10 min. The chromatin was sheared using a Bioruptor (Diagenode, high power, 20 cycles of 30 s with 30 s recovery) to obtain a mean fragment size of ~300 bp. Insoluble material was removed by centrifugation. The chromatin solution was pre-cleared with protein A sepharose (Sigma-Aldrich), 10% retained (input) and then chromatin immunoprecipitated overnight using 5 μg primary antibody (rabbit IgG or rabbit α-TASOR) and protein A sepharose. The next day the beads were washed a total of five times, and then bound protein-DNA complexes eluted in 0.15 M NaHCO_3_ and 1% SDS. Cross-links were reversed by overnight incubation at 67°C with 0.3 M NaCl and 1 μg RNase A. Proteinase K (60 μg) was then added and the samples incubated for 2 h at 45°C. DNA was purified using a spin column (Qiagen PCR purification kit). Illumina sequencing libraries were produced from this material using the TruSeq kit (Illumina), and sequenced on a HiSeq 2500 instrument. Single-end reads (1×50) were aligned to the human genome (hg38) using Bowtie2 with default parameters and converted to bam files with samtools (Langmead and Salzberg, 2012; Li et al., 2009b). Coverage plots for input and ChIP (IgG and TASOR) samples were generated using bamCoverage (deepTools), with reads extended (250 bp) and normalized using RPGC (reads per genomic context; chrX ignored) with an effective genome size of 2913022398 (hg38) (Ramírez et al., 2014). ChIP experiments were done once. Normalized bigwig files were displayed in IGV (Robinson et al., 2011) and heatmaps generated with deepTools computeMatrix and plotHeatmap commands (Ramírez et al., 2014).

### CUT&Tag

We followed the protocol detailed by the Henikoff lab (Kaya-Okur et al., 2019) with alterations made after consultation with protocol authors, or due to the method being under regular review and optimization. 100,000 cells were washed twice (20 mM HEPES pH 7.5, 0.15 M NaCl, 0.5 mM spermidine, 1x Roche complete protease inhibitors) and attached to activated ConA-coated magnetic beads (Bangs Laboratories) at RT for 15 min. Cells bound to the beads were resuspended in 100 μL buffer (20 mM HEPES pH 7.5, 0.15 M NaCl, 0.5 mM Spermidine, 1x Roche complete protease inhibitors, 0.05% digitonin (Millipore), 2 mM EDTA) containing primary antibody (1:50 dilution). Incubation proceeded at RT for 2 h with gentle shaking. Tubes were placed on a magnet stand to allow removal of unbound antibody. The secondary antibody (guinea pig anti-rabbit IgG, 0.25 g/L) was added at 1:100 dilution and cells incubated at RT for 1 h with gentle shaking. Cells were washed three times on the magnet in 1 mL buffer (20 mM HEPES pH 7.5, 150 mM NaCl, 0.5 mM Spermidine, 1x Roche complete protease inhibitors, 0.05% digitonin). Meanwhile pA-Tn5 adapter complex (40 nM, a generous gift from Steve Henikoff) was prepared in a higher salt, lower digitonin buffer (20 mM HEPES, pH 7.5, 0.35 M NaCl, 0.5 mM spermidine, 1x Roche complete protease inhibitors, 0.01% digitonin). This buffer had a slightly increased NaCl concentration over that recommended in the protocol, to reduce non-specific binding to open chromatin (the so-called ATAC-seq artifact). After the final wash, 100 μL of pA-Tn5 solution was added to the bead-bound cells with gentle vortexing and the cells incubated at RT for 1 h with gentle shaking. Cells were then washed three further times in 1 mL buffer, before resuspension in 50 μL tagmentation buffer (20 mM HEPES pH 7.5, 0.35 M NaCl, 10 mM MgCl_2_, 0.5 mM spermidine, 1x Roche complete protease inhibitors, 0.01% digitonin). Tagmentation was allowed to proceed at 37 °C for 1 h before quenching with 20 mM EDTA, 0.5% SDS (both final concentrations) and 10 μg Proteinase K (Thermo Fisher Scientific). The mixture was incubated at 37 °C overnight. The next day, tubes were incubated at 70 °C for 20 min to further inactivate the protease. DNA was extracted from the mixture with 2.2x SPRI beads (KAPA). After twice washing the beads with 80% EtOH, DNA was eluted in 25μL water. For PCR, 21 μL DNA was mixed with 2 μL of 10 μM universal i5 primer (AATGATACGGCGACCACCGAGATCTACACTCGTCGGCAGCGTCAGATGTG) and 2 μL of 10 μM uniquely-barcoded i7 primer (CAAGCAGAAGACGGCATACGAGAT[i7]GTCTCGTGGGCTCGGAGATGT), then 25 μL NEBNext HiFi 2x PCR Master mix was added and pipette mixed. The following thermocycler program was used: 72 °C for 5 min; 98 °C for 30 s; 13 cycles of 98 °C for 10 s and 63 °C for 30 s; final extension at 72 °C for 1 min and hold at 8 °C. Post-PCR clean-up was performed with 1.1x SPRI beads (KAPA). Libraries were pooled in approximately equimolar ratios based on capillary electrophoresis (Agilent) and/or fluorometry (Thermo Fisher Scientific) quantification results, before final left-sided size selection (1.1x SPRI) to remove residual PCR primers. Paired end reads (2×150 bp) were generated on a HiSeq 4000 instrument (Illumina). Reads were aligned to the human genome (hg38) using Bowtie2 (--local –very-sensitive-local –no-mixed –no-discordant -I 10 -× 700) and converted to bam files with samtools (Langmead and Salzberg, 2012; Li et al., 2009b). PCR duplicates were removed using Picard (http://broadinstitute.github.io/picard/) before conversion to bedgraph file format (bedtools (Quinlan and Hall, 2010)). Coverage plots for display and comparison of tracks were generated using bamCoverage (deepTools (Ramírez et al., 2014)) after downsampling bam files to the control/KO sample based on the final number of mapped reads. We note that this is a conservative approach, because library complexity is related to the number of true binding sites in such a targeted experiment (Steve Henikoff, personal communication). CUT&Tag experiments to assess TASOR binding were done in biological duplicate. Resulting bigwig files were displayed in IGV (Robinson et al., 2011) and heatmaps made with deepTools computeMatrix and plotHeatmap packages (Ramírez et al., 2014).

### RNA-seq analysis

Paired end (2×150) reads from HeLa RNA-seq experiments that we reported elsewhere (Tchasovnikarova et al., 2017) were re-mapped to hg38 using HiSat2 (--no-mixed –no-discordant) then converted to bam files in samtools (Kim et al., 2019; Li et al., 2009b). Fragments overlapping representative transcripts from annotated human genes (gencode v29) were counted in the featureCounts tool (--primary –fraction -t exon -p) from the subread package (Liao et al., 2014) and the mean taken of the three replicates (n=3).

### BioID

The following protocol was adapted from (Schopp et al., 2017). TASOR knockout cells expressing HA-BirA-tagged TASOR variants were grown in square 500 cm^2^ dishes (Corning) in DMEM, which was supplemented with 50 μM biotin for 18 h prior to downstream processing. After washing with PBS, cells were scraped in PBS before pelleting (400g, 5 min, RT). Nuclei were isolated by resuspending cells in 10 mL nuclear isolation buffer (1 mM HEPES, 85 mM KCl, 0.5% IGEPAL) before being pelleted again (800g, 5 min, 4 °C). Nuclear pellets were lysed by sonication on ice in 10 mL lysis buffer (50 mM Tris pH 7.4, 0.5 M NaCl, 0.4% SDS, 5 mM EDTA, 1 mM DTT, 1x Roche complete protease inhibitor). After sonication, Triton X-100 concentration was adjusted to 2% and NaCl concentration to 150 mM. Lysates were then centrifuged (16,000 g, 10 min, 4 °C). Protein concentration was measured by BCA assay (Pierce) and equal amounts taken for further steps. Samples were incubated with 50 μL pre-equilibrated magnetic Dynabeads MyOne Streptavidin C1 at 4 °C overnight on a rotating wheel. The next day, a series of washing steps were carried out at RT, each twice for 5 min unless otherwise stated. Wash buffer 1 contained 2% SDS, 50 mM TEAB pH 8.5, 10 mM TCEP, 20 mM iodoacetamide (30 min incubation). Wash buffer 2 contained 50 mM HEPES pH 7.4, 1 mM EDTA, 500 mM NaCl, 1% Triton X-100, 0.1% Na-deoxycholate. Wash buffer 3 contained 10 mM Tris pH 8.0, 0.25 M LiCl, 1 mM EDTA, 0.5% NP-40, 0.5% Na-deoxycholate. Wash buffer 4 contained 50 mM Tris pH 7.4, 50 mM NaCl, 0.1% NP-40. Wash buffer 5 contained 50 mM TEAB pH 8.5, 6 M urea. Wash buffer 6 contained 50 mM TEAB pH 8.5. Finally, beads were resuspended in 50 μL 50 mM TEAB pH 8.5 containing 0.1% Na-deoxycholate and 200 ng of trypsin (Promega), before incubation overnight at 37 °C with periodic shaking (Eppendorf Thermomixer). After removing the beads digests were subjected to clean-up using SDP-RPS extraction material (Affinisep) backed into 200 μL pipette tips. Columns were conditioned with 100 μL ACN followed by 100 μL 0.5% TFA. Samples were acidified with a final concentration of 0.5% TFA and an equal volume of ethyl acetate loaded onto the columns. Columns were then washed with 100 μL 0.2% TFA and 100 μL ethyl acetate and eluted in 80% ACN + 5% ammonium hydroxide. Samples were dried under vacuum and stored at −20 °C prior to analysis.

### Mass spectrometry

Samples were resuspended in 10 μL 5% DMSO, 0.5% TFA and the whole sample injected. Data were acquired on an Orbitrap Fusion mass spectrometer (Thermo Scientific) coupled to an Ultimate 3000 RSLC nano UHPLC system (Thermo Scientific). Samples were loaded at 10 μL/min for 5 min on to an Acclaim PepMap C18 cartridge trap column (300 μm × 5 mm, 5 μm particle size) in 0.1% TFA. After loading, a linear gradient of 3-32% solvent B over 60 min was used for sample separation with a column of the same stationary phase (75 μm × 75 cm, 2 μm particle size) before washing at 90% B and re-equilibration. Solvents were A: 0.1% FA and B: ACN/0.1% FA. MS settings were as follows. MS1: quadrupole isolation, 120 000 resolution, 5e5 AGC target, 50 ms maximum injection time, ions accumulated for all parallelisable time. MS2: quadrupole isolation at an isolation width of m/z 0.7, HCD fragmentation (NCE 34) with the ion trap scanning out in rapid mode from, 8e3 AGC target, 0.25 s maximum injection time, ions accumulated for all parallelisable time. Target cycle time was 2 s. Spectra were searched by Mascot within Proteome Discoverer 2.2 in two rounds of searching. The first search was against the Uniprot human reference proteome and compendium of common contaminants (GPM). The second search took all unmatched spectra from the first search and searched against the human trEMBL database. The following search parameters were used. MS1 Tol: 10 ppm, MS2 Tol: 0.6 Da, fixed mods: carbamindomethyl (C); var mods: oxidation (M), enzyme: trypsin (/P). Peptide spectrum match (PSM) FDR was calculated using Mascot percolator and was controlled at 0.01% for ‘high’ confidence PSMs and 0.05% for ‘medium’ confidence PSMs. Proteins were quantified using the Minora feature detector within Proteome Discoverer.

### Western blotting

Cells were lysed in 1% SDS plus 1:100 (v/v) benzonase (Sigma) for 15 min at room temperature, and then heated to 65 °C in SDS sample loading buffer for 5 min. Following separation by SDS-PAGE, proteins were transferred to a PVDF membrane (Millipore), which was then blocked in 5% milk in PBS + 0.2% Tween-20. Membranes were probed overnight with the indicated primary antibodies, washed four times in PBS + 0.2% Tween-20, then incubated with HRP-conjugated secondary antibodies for 1 h at RT. Reactive bands were visualized using SuperSignal West Pico (Thermo Fisher Scientific). Alternatively, after SDS-PAGE, proteins were transferred to a nitrocellulose membrane (Thermo Fisher Scientific iBlot2), which was blocked in 5% milk in PBS (no detergent) for 1h. Membranes were probed overnight with the indicated primary antibodies in 5% milk in PBS + 0.1% Tween-20, washed thoroughly in PBS + 0.1% Tween-20, then incubated with DyLight-680 or 800-conjugated secondary antibodies (Thermo Fisher Scientific) at 1:10,000 dilution for 30 min at RT. After thorough washing with PBS-Tween, PBS and then water, blots were imaged on the Odyssey near-infrared system (LI-COR).

### Subcellular fractionation

Cells were washed twice in PBS and once in buffer A (10 mM HEPES pH 7.9, 1.5 mM MgCl_2_, 10 mM KCl, 0.5 mM dithiothreitol (DTT) and protease inhibitor cocktail). Cells were then pelleted and resuspended in buffer A with 0.1% (v/v) NP40 and incubated on ice for 10 min. The supernatant containing the cytoplasmic fraction was collected following centrifugation (1,300 g, 4 min, 4°C) and further clarified by high-speed centrifugation (20,000 g, 15 min, 4 °C). The remaining pellet was washed in buffer A without NP40 and resuspended in an equal volume (relative to the cytoplasmic extract) of buffer B (20 mM HEPES pH 7.9, 1.5 mM MgCl_2_, 0.3 M NaCl, 0.5 mM DTT, 25% (v/v) glycerol, 0.25% Triton X-100, 0.2 mM EDTA and protease inhibitor cocktail). The supernatant containing the soluble nuclear fraction was collected following centrifugation (1,700 g, 4 min, 4 °C), and the insoluble pellet, composed primarily of chromatin and associated proteins, was resuspended in an equal volume of Laemmli buffer (relative to the cytoplasmic and soluble nuclear extracts). Equal volumes of cytoplasmic, soluble and insoluble nuclear fractions were separated by SDS–PAGE, transferred to a PVDF membrane (Millipore) and probed with relevant antibodies.

### Confocal immunofluorescence microscopy

Cells were fixed for 15 minutes with 4% formaldehyde in PBS, permeabilized for 5 minutes with 0.1% TritonX-100 in PBS and then blocked with 2% BSA in PBS for 1 hour. Cells were stained with primary antibodies at 1:200 dilution in blocking buffer and, after further washing, with secondary antibody (anti-rabbit AlexaFluor 647, 1:500 dilution) in blocking buffer for 1 h. Samples were washed thoroughly and cover slips mounted on microscopy glasses with ProLong Gold anti-fade reagent with DAPI (Invitrogen). Imaging was performed using Nikon Ti microscope equipped with CSU-X1 spinning disc confocal head (Yokogawa) and with Zeiss 780 system.

### Protein expression and purification

A synthetic *E. coli* codon-optimized DNA construct (IDT) encoding TASOR residues 106-332 was cloned into the expression vector pET-15b for production of the N-terminally thrombin-cleavable His_6_-tagged protein product (MGSSHHHHHHSSGLVPRGSHM[…]). Mutation of this construct to generate variants Y305A or the construct used for crystallography (110-332 Δ261-269) were done with standard methods. Transformed *E. coli* BL21(DE3) cells (NEB) were grown at 37 °C in 2xTY media containing 100 mg/L ampicillin. Expression was induced at an OD_600_ of 0.8 with 0.2 mM IPTG for 18 h at 18 °C. The culture was pelleted and resuspended in a buffer containing 50 mM Tris pH 8.0, 0.15 M NaCl, 10 mM imidazole, 1 mM DTT and 1x Roche complete EDTA-free protease inhibitors, then flash frozen in liquid nitrogen and stored at −80 °C. All subsequent steps were done at 4 °C unless otherwise stated. Further lysis was achieved by extensive sonication (3 × 3 min). A solution of benzonase (1:10,000 v/v final concentration, Sigma) was added and after 30 min incubation with stirring, the NaCl concentration was adjusted to 0.5 M, otherwise the protein co-purified with host RNA. The lysate was clarified by centrifugation (15,000 g, 45 min) and the protein-containing supernatant bound to preequilibrated Ni-NTA beads (Generon) for 1 h with rocking. The beads were washed with at least 20 CV Ni wash buffer (50 mM Tris pH 8.0, 0.5 M NaCl, 10 mM imidazole, 1 mM DTT) before a stepwise elution in batch mode with 3 × 5 CV of Ni wash buffer supplemented with 0.2 M, 0.3 M and 0.5 M imidazole. Further purification was achieved with size-exclusion chromatography on a Superdex 200 increase (10/300) column (GE) in buffer (50 mM HEPES pH 7.5, 0.2 M NaCl, 0.5 mM TCEP). For crystallography trials and NMR experiments, the His_6_-tag was cleaved with restriction-grade thrombin (Millipore) overnight on ice in Tris buffer supplemented with 2.5 mM CaCl_2_. The next day the protease was removed by incubation with 100 μL benzamidine sepharose beads (GE) for 5 min at RT. For crystallography trials an additional ion exchange chromatography step with a monoS column was included in the protocol. Tag cleavage was done after the Ni-affinity step, before size exclusion and ion exchange chromatography steps. For SeMet-labelled protein, expression was repeated in minimal media containing L-(+)-selenomethionine (Anatrace), using an established strategy (Van Duyne et al., 1993). Single ^15^N labelling, double ^13^C/^15^N labelling or triple ^2^H/^13^C/^15^N labelling for NMR experiments required expression in minimal media made with ^15^NH_4_Cl, ^13^C glucose and/or D_2_O (Sigma) as appropriate. Cultures in D_2_O grew more slowly and were therefore kept at 25 °C throughout. Purification of these labelled samples otherwise followed the protocol used for unlabeled samples.

For co-expression of the TASOR-MPP8 complex in *E. coli*, we co-transformed BL21(DE3) cells with the pET15b vector harbouring His-tagged TASOR(354-633) and the pRSF vector harbouring MPP8(527-860). The proteins were expressed by growing the cells in autoinduction media for 60 h at 18 °C, and the complex purified by Ni-affinity purification. Eluates were concentrated and analysed by SEC-MALS at 293 K using a Superdex 200 (10/300) column in a buffer containing 20 mM HEPES pH 7.5, 0.5 M NaCl, 0.5 mM TCEP. Light scattering analysis was performed in the ASTRA software package (Wyatt), using band broadening parameters obtained from a BSA standard run on the same day under identical conditions. MALS data were used to fit the average molar mass across the complex peak (quoted to the nearest kDa).

Expression of full-length His-tagged *Hs*PARP-1 and the in vitro gel-based PARylation assay followed a detailed protocol published elsewhere (Langelier et al., 2011).

### NMR

NMR data were collected at 298 K using a Bruker Avance II+ 700 MHz spectrometer with triple resonance cryoprobe unless otherwise stated. All samples were prepared with 5% D_2_O as a lock solvent, in PBS (pH 7.0) supplemented with 1 mM TCEP and 0.05% w/v NaN_3_, and degassed prior to data acquisition. ^1^H-^15^N BEST-TROSY (band selective excitation short transient transverse relaxation optimized spectroscopy) spectra were collected for the ^15^N labelled WT TASOR sample and the ^2^H/^13^C/^15^N WT TASOR sample using an optimized pulse sequence (Favier and Brutscher, 2011). An initial, incomplete assignment of WT TASOR was carried out using standard TROSY based triple resonance spectra with deuterium decoupling: trHNCO and trHNCACO with 2048*64*128 complex points in the ^1^H, ^15^N and ^13^C dimensions respectively, trHNCA and trHNCOCA with 2018*64*160 complex points in the ^1^H, ^15^N and ^13^C dimensions respectively and trHNCACB and trHNCOCACB with 2048*64*110 complex points in the ^1^H, ^15^N and ^13^C dimensions respectively. This assignment revealed a subset of residues without peak data. A comparison of 2D projections with a limited number of equivalent triple resonance experiments collected on a 15N/13C only labelled sample revealed additional peak data. This indicated that the deuterated sample had incomplete back exchange of the solvent exchangeable backbone NH protons within the core of the protein. Partial denaturation of the ^13^C/^15^N/^2^H sample with 3.5 M urea in PBS, followed by incremental stepwise dilution of urea back to 0 M, allowed back-exchange and additional data sets were collected to complete the assignment. These additional experiments included trHNCACB, trHNCA, trHNCACO, trHNCO and trHNCOCA spectra recorded as above. All triple resonance data sets were collected with 20-40% Non-Uniform Sampling (NUS) and processed using compressed sensing (Kazimierczuk and Orekhov, 2011). All 2D datasets were processed using Topspin version 3.1 or higher (Bruker) and all spectra analyzed using Sparky 3.115. The assignment was completed for 184/221 non-proline backbone amide resonances using MARS (Jung and Zweckstetter, 2004). The dynamic properties of TASOR were investigated using standard Bruker T_1_ and T_2_ relaxation pulse programs. T_1_ and T_2_ data sets were collected on an Avance III HD 800 MHz spectrometer fitted with a triple resonance cryoprobe and T_1_ delays of 50, 100, 200, 500, 800, 1500, 2200 and 3000 ms and T_2_ delays of 16, 32, 48, 64, 96, 128, 160 and 192 ms. Signal intensity measurements and slope fitting was completed using Sparky. A second, higher-resolution T_2_ data set was collected at 950 MHz (Bruker Avance III HD) using the same relaxation delays.

### X-ray crystallography

TASOR PARP domain (residues 110-332 with internal deletion of residues 261-269 and His_6_ tag cleaved), native or SeMet-labelled, was concentrated to 8 g/L (320 μM) in buffer containing 20 mM HEPES pH 7.5, 0.15 M NaCl, 0.5 mM TCEP. Crystals were grown at 18 °C by the sitting drop vapor diffusion method, by mixing the protein at a 1:1 ratio with reservoir solution containing 0.1 M MES pH 6.5, 0.1 M NaP_i,_, 0.1 M KP_i_, 2 M NaCl. Crystals appeared in overnight and were frozen within 2 d in liquid N_2_ using paraffin oil as a cryoprotectant. X-ray diffraction data were collected at 100 K at Diamond Light Source beamlines i02 and i03 and processed using autoproc or xia2 packages (Vonrhein et al., 2011; Winter, 2010). Automatic experimental phasing was done in AutoSol (Phenix (Adams et al., 2010)) using the single anomalous dispersion (SAD) method with selenium as the heavy atom. The resulting model was built and refined in Coot (Emsley and Cowtan, 2004) and Phenix, before being used as a search model for the native dataset in Phaser (McCoy et al., 2007). Further model building and refinement were done with this dataset in Coot and Phenix. Crystallographic data are summarized in Table S2.

### Fluorescence polarization

TASOR(106-332) was titrated into RNA-binding buffer (20 mM HEPES pH 7.5, 50 mM NaCl, 2 mM MgCl_2_, 0.5 mM TCEP) containing 33 nM of a single-stranded RNA oligonucleotides 5’-labelled with 6-carboxyfluorescein (6-FAM). Fluorescence polarization of 30-μl samples was measured in 384-well black, clear-bottomed plates (Corning) with a ClarioSTAR plate reader using 482/530 nm filters.

### Differential scanning fluorimetry (DSF)

10 μL samples containing 10 μM protein (TASOR variants or PARP1 catalytic domain) in the presence or absence of benzamide (Sigma) were loaded into glass capillaries (Nanotemper) by capillary action. Intrinsic protein fluorescence at 330 nm and 350 nm was monitored between 15 and 90 °C in the Prometheus NT.48 instrument (Nanotemper), and the T_m_ values calculated within the accompanying software by taking the turning point of the first derivative of the F350:F330 ratio as a function of temperature.

## Data availability

The NMR data were deposited in the Biological Magnetic Resonance Bank, dataset ID 50094. The structure factors and atomic coordinates for the crystal structure were deposited in the Protein Data Bank with code PDB: 6TL1. The original experimental X-ray diffraction images were deposited in the SBGrid Data Bank (data.SBGrid.org), with Data ID 742, DOI:10.15785/SBGRID/742. The CUT&RUN, ChIP and CUT&Tag data were deposited in XXX, ID XXX.

## Acknowledgements

We thank Stuart Bloor, Thomas Monteiro-Crozier, Ana Teixera-Silva, Frank Adolf and other members of the Lehner and Modis labs for useful discussions. We thank Steve Henikoff and members of the Henikoff group (Fred Hutchinson Cancer Center) for advice and generous support on CUT&RUN and CUT&Tag protocols through protocols.io, and for kindly supplying purified pA-MNase and pA-Tn5 reagents. We thank Tom Odgen and David Neuhaus for advice on full-length PARP1 purification and for the gift of an aliquot of PARP1 catalytic domain. NMR studies were supported by the Francis Crick Institute (FCI) through provision of access to the MRC Biomedical NMR Centre. The FCI receives its core funding from Cancer Research UK (FC001029), the MRC (FC001029), and the Wellcome Trust (FC001029). Crystallographic data were collected on beamline I02 and I03 at Diamond Light Source (DLS). Access to DLS (proposal MX11235) was supported by the Wellcome Trust, MRC and BBSRC. We thank Minmin Yu for advice on crystallographic data collection strategies. We thank Toby Darling and LMB Scientific Computing for IT support behind genomics analysis. I.A.T. is a fellow of the Damon Runyon Cancer Research Foundation and R.T.T. is a Sir Henry Wellcome Postdoctoral Fellow (201387/Z/16/Z). This work was supported by Wellcome Trust Senior Research Fellowships 101908/Z/13/Z and 217191/Z/19/Z to Y.M., a Wellcome Trust Principal Research Fellowship 101835/Z/13/Z to P.J.L and a BBSRC Future Leader Fellowship BB/N011791/1 to C.H.D.

## Author Contributions

Conceptualization, C.H.D., I.A.T., R.T.T., D.M.P., P.J.L. and Y.M.; Methodology, C.H.D., I.A.T., R.T.T, A.V.P., M.S., A.A., J.C.W, S.M.V.F., P.J.L. and Y.M.; Investigation, C.H.D., I.A.T., R.T.T., A.V.P., D.M.P., A.A. and S.M.V.F.; Validation – genomics, C.H.D., I.A.T., R.T.T., and A.V.P.; Validation – NMR, C.H.D., J.W. and S.M.V.F; Validation – crystal structure, C.H.D. and Y.M.; Writing – Original Draft, C.H.D., I.A.T. and R.T.T.; Writing – Review & Editing, C.H.D, I.A.T., R.T.T., P.J.L and Y.M.; Visualization, C.H.D., I.A.T., R.T.T., A.V.P., A.A. and J.W.; Supervision, P.J.L. and Y.M.; Project Administration, C.H.D, P.J.L and Y.M.; Funding Acquisition, C.H.D, P.J.L. and Y.M.

## Supplementary Figures and Tables

**Fig. S1.**
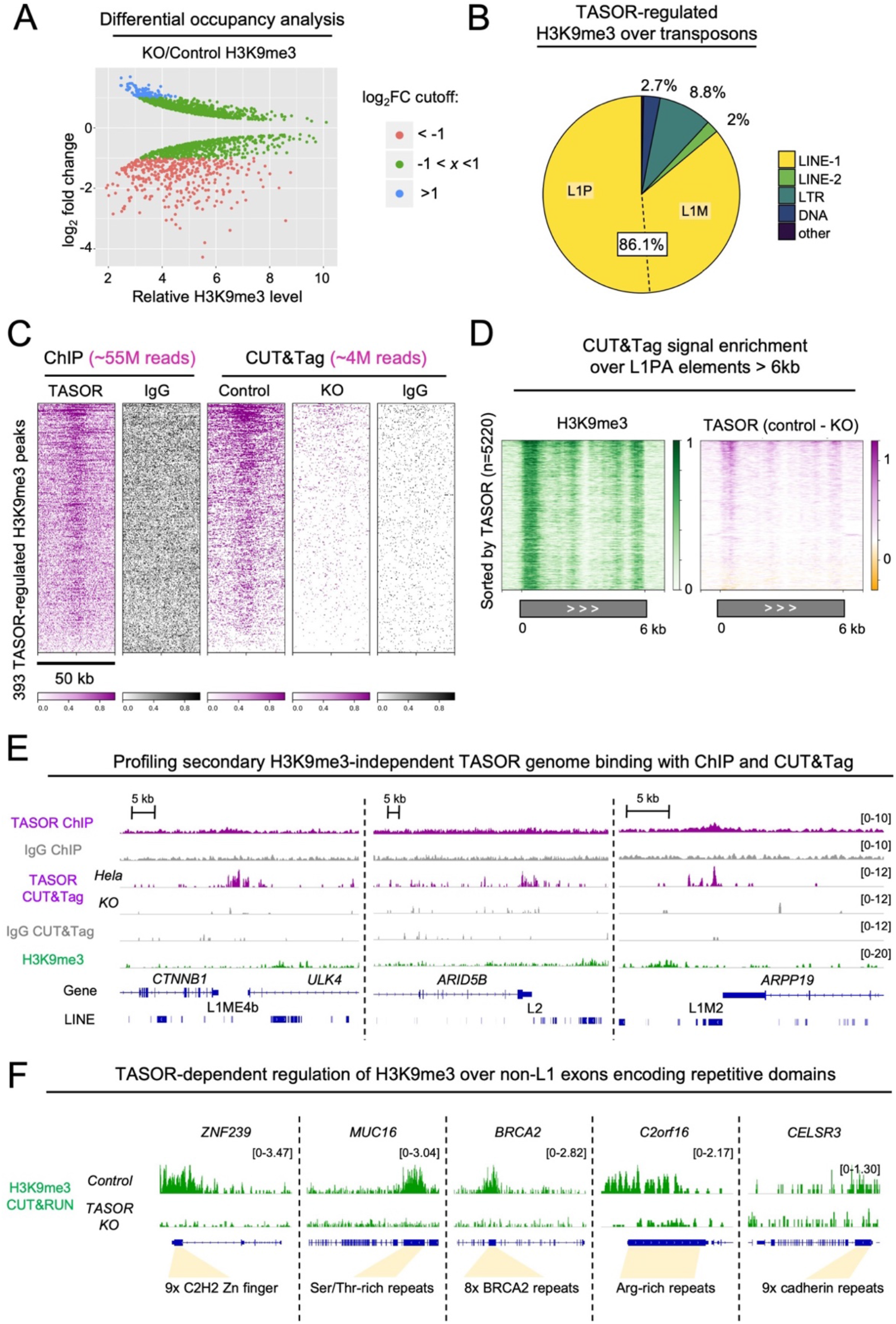
(**A**) A log_2_-fold change <-1 of normalized H3K9me3 counts for TASOR-negative cells versus TASOR-positive cells, and FDR <0.05, was used to define 393 TASOR-regulated sites. (**B**) Overlaps of 393 TASOR-regulated H3K9me3 peaks with different repeat classes. Since these peaks extend over several kilobases (mean length 6369 nt), several peaks covered more than one repeat annotation. (**C**) Heatmaps of the signal from ChIP-TASOR, ChIP-IgG, CUT&Tag-TASOR and CUT&Tag control experiments across 393 TASOR-regulated loci (rows). (**D**) CUT&Tag H3K9me3 and TASOR signal plotted over 5,220 L1PA elements longer than 6 kb in the hg38 assembly. (**E**) Snapshots showing three loci with evidence of H3K9me3-independent binding by TASOR using ChIP and CUT&Tag. (**F**) Genome browser snapshots showing TASOR-dependent H3K9me3 regulation over repetitive exons that were not LINE-1 elements.

**Fig. S2.**
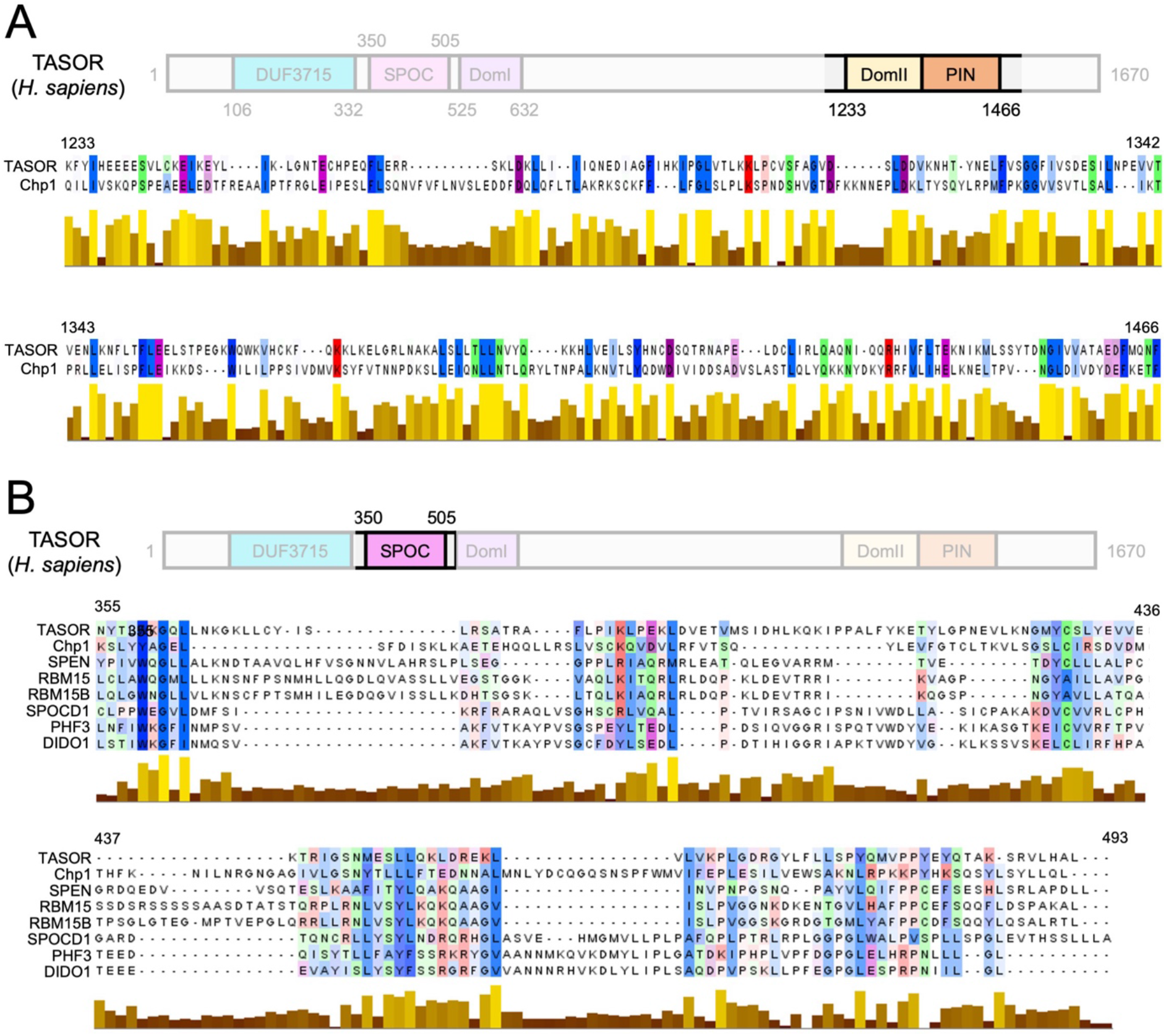
(**A**) Sequence alignment of the DomII/PIN domains from TASOR and *S. pombe* Chp1. (**B**) Sequence alignment of SPOC domains from TASOR, Chp1 and six other SPOC-containing proteins.

**Fig. S3.**
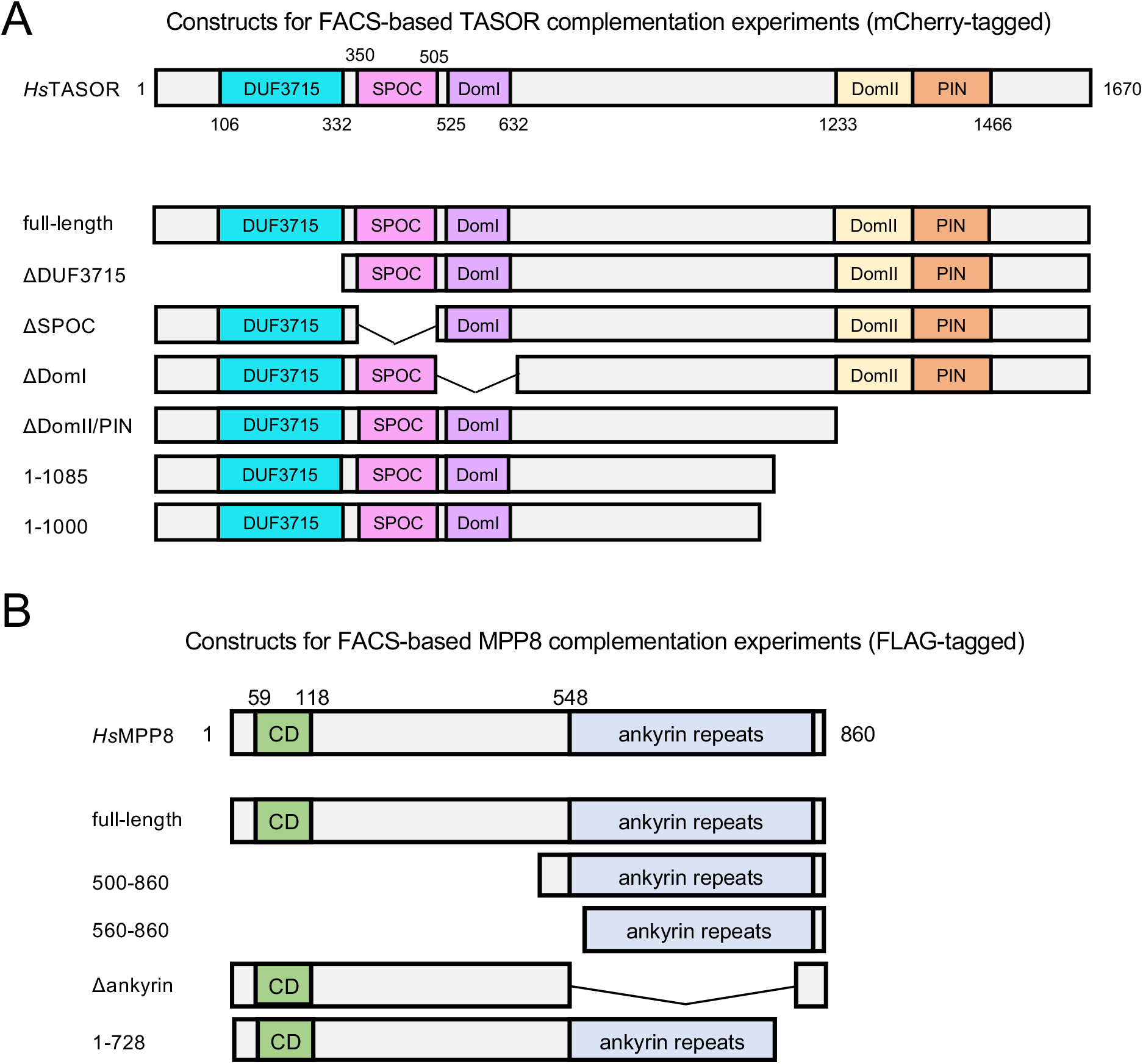
(**A**) Summary of TASOR domain deletion constructs for FACS-based transgene repression assays. (**B**) Summary of MPP8 domain deletion constructs for FACS-based transgene repression assays.

**Fig. S4.**
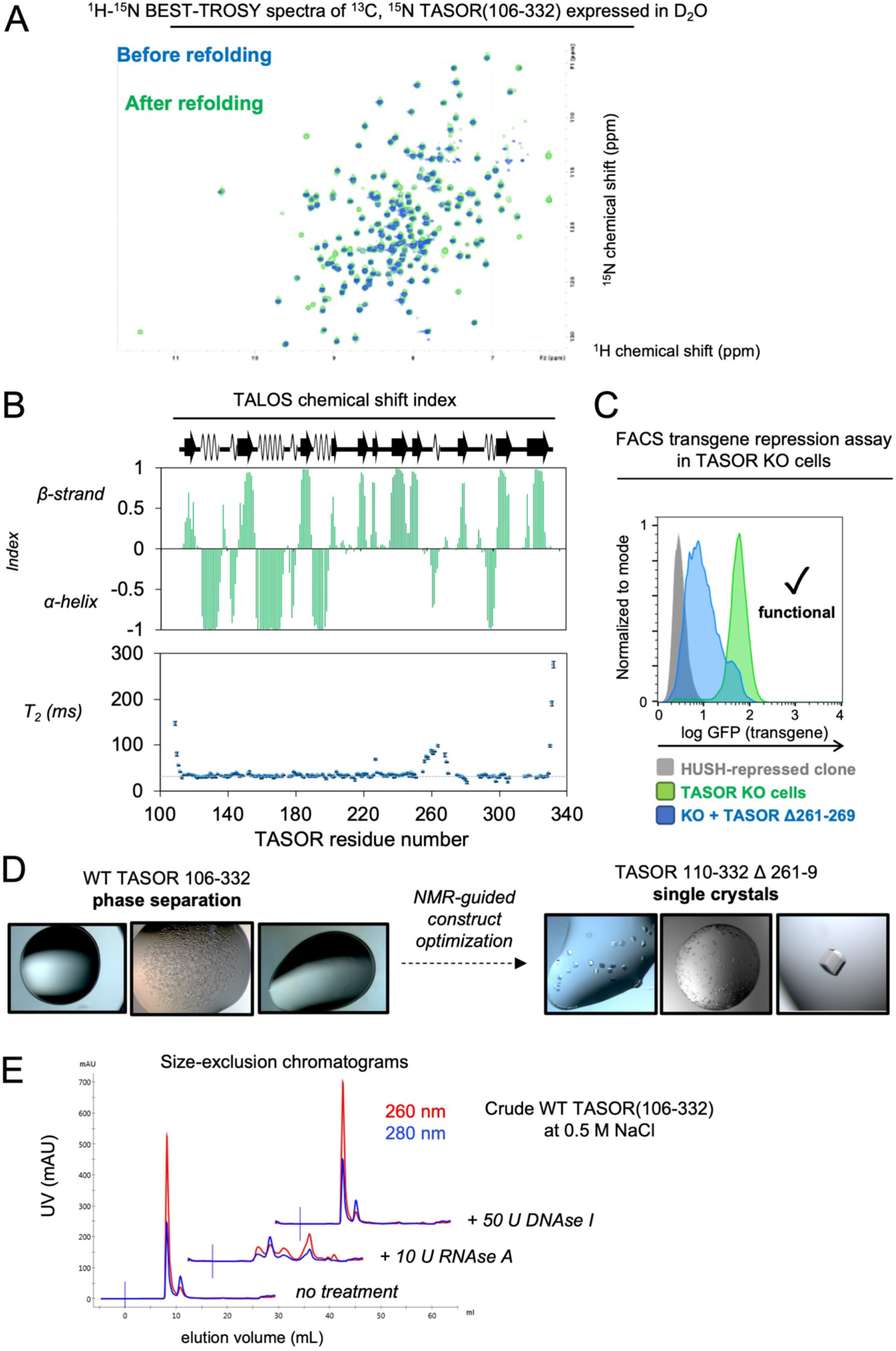
(**A**) NMR spectra showing denaturation/refolding experiment to enforce H/D exchange of core amide deuterons. (**B**) Chemical shift indexing of TASOR DUF3715 (residues 106-332). (**C**) FACS transgene repression assay showing 261-269 loop deletion does not affect TASOR function. (**D**) Optimization of DUF3715 construct for crystallization experiments. (**E**) Crude TASOR DUF3715 bound to expression host RNA but not DNA following Ni-affinity purification. Shown are S200 SEC traces following the treatments as annotated, under otherwise identical conditions.

**Fig. S5.**
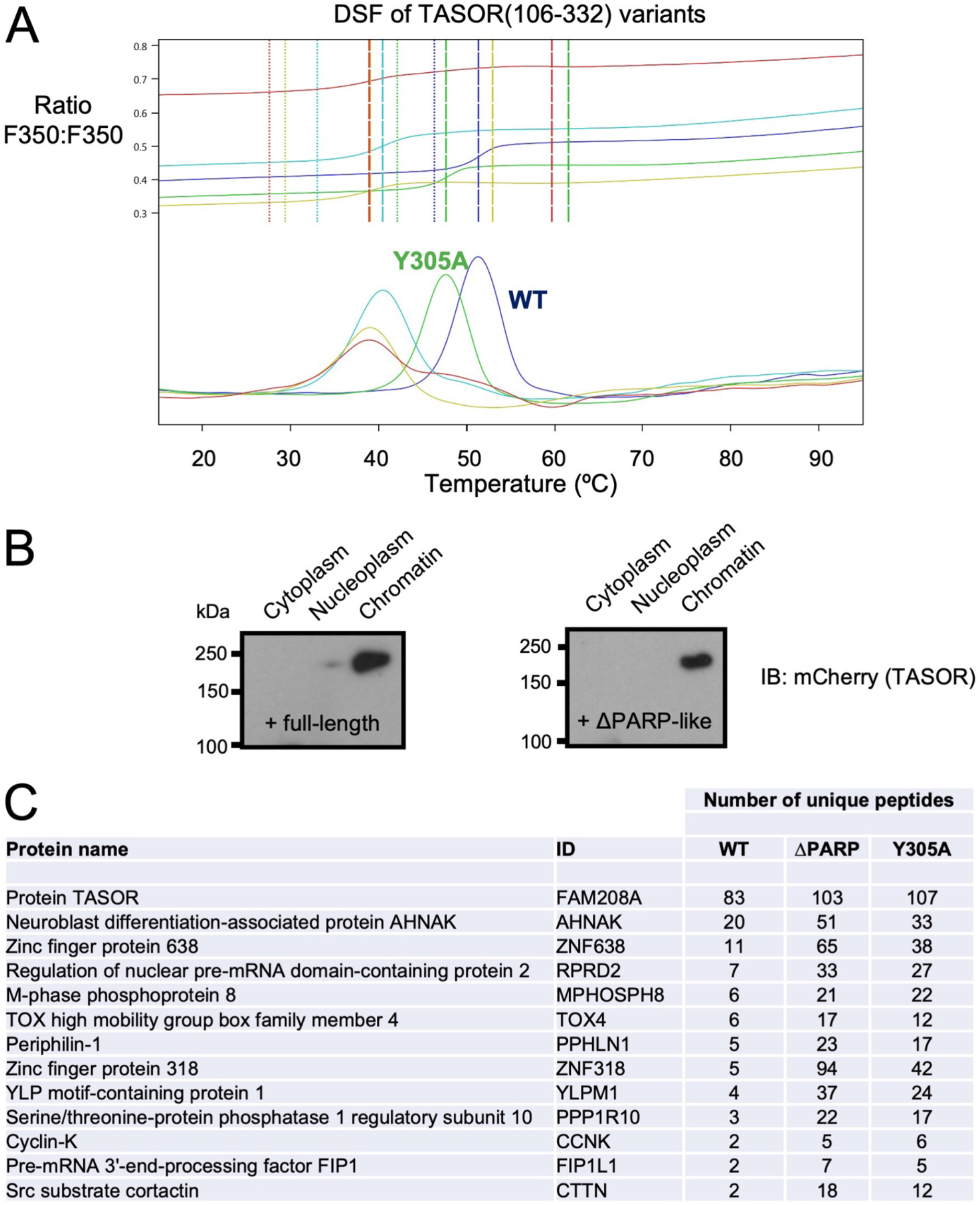
(**A**) Differential scanning fluorimetry of WT and Y305A TASOR(106-332) shows both proteins are folded. The upper panel shows the ratio of fluorescence at 350 nm and 330 nm (excitation: 280 nm); the lower panel shows the first derivative of this ratio as a function of temperature (*x* axis). A turning point in the derivative trace defines the melting temperature (T_m_) of the domain. (**B**) Subcellular fractionation of full-length TASOR and a PARP deletion construct. (**C**) BioID hits for full-length (WT), PARP deletion and Y305A TASOR variants. Listed are those hits with more than two unique peptides in the WT experiment.

**Table S1.** Crystallographic data collection and refinement statistics for TASOR pseudo-PARP.

|  | Native | Se-SAD |
| --- | --- | --- |
| **Data collection** |  |  |
| X-ray source | DLS i02 | DLS i03 |
| Space group | *P4<sub>1</sub>2<sub>1</sub>2* | *P4<sub>1</sub>2<sub>1</sub>2* |
| Cell dimensions |  |  |
| a = b, c (Å) | 74.59, 184.0 | 74.45, 182.3 |
| α = β = γ (°) | 90 | 90 |
| Wavelength (Å) | 0.97949 | 0.97980 |
| Resolution (Å) | 57.94 – 2.03 (2.08 – 2.03)* | 91.14 – 2.27 (2.39 – 2.27)* |
| Observations | 438905 | 350588 |
| Unique reflections | 34589 | 24704 |
| *R*<sub>merge</sub> | 0.094 (1.460) | 0.115 (1.296) |
| <*I*> / σ(*I*) | 16.1 (2.1) | 18.4 (2.5) |
| Completeness (%) | 100.0 (100.0) | 100.0 (100.0) |
| Redundancy | 12.7 (12.6) | 14.2 (14.4) |
| CC<sub>ano</sub> | – | 0.846 (0.008) |
| **Refinement** |  |  |
| Protein molecules in a.u. | 2 |  |
| *R*<sub>work</sub> / *R*<sub>free</sub> | 0.195 / 0.229 |  |
| No. of non-H atoms |  |  |
| Protein | 3359 |  |
| Solvent | 146 |  |
| Mean B-factors |  |  |
| Protein | 56.6 |  |
| Solvent | 49.3 |  |
| R.m.s deviations |  |  |
| Bond lengths (Å) | 0.007 |  |
| Bond angles (°) | 1.02 |  |
| Ramachandran statistics |  |  |
| % favored | 98.0 |  |
| % allowed | 2.0 |  |
| % outliers | 0.0 |  |

\*Highest resolution shell is shown in parentheses.
A.u, asymmetric unit; R.m.s., root mean square.

